# Myeloid Lectin Profiling Identifies SYK as a Targetable Signaling Node for Remodeling Immunosuppressive Tumor-Associated Macrophages in Breast Cancer

**DOI:** 10.64898/2026.09.04.749378

**Authors:** Gonçalo Trindade, Giacomo Domenici, Miguel Pinto, Viviana Correia, Sofia Batalha, Nádia Duarte, Angelina Sá Palma, Inês A. Isidro, Catarina Brito

**Author notes:** Corresponding Author: Apartado 12, 2781-901 Oeiras, Portugal.

## Abstract

Immunosuppressive tumor microenvironments (TMEs) are common in breast cancer (BC), where tumor-promoting myeloid-driven inflammation contributes to immune dysfunction and poor immunotherapy response. Lectins expressed by tumor-associated macrophages (TAMs) act as glycan-sensing immunoregulatory receptors driving immunosuppression, but whether specific myeloid lectins define immunosuppressive TAM states, or their signaling is pharmacologically targetable remain unclear. We used an orthogonal prioritization approach combining immunosuppressive stratification of the SCAN-B cohort (3207 patients), differential expression, single-cell myeloid localization, and immune-related upstream-regulator inference. Immunosuppressive tumors were enriched for Basal-like and HER2-enriched subtypes, with worse survival. This identified a 12-lectin panel, which was probed in a human 3D immunosuppressive TME model combining BC spheroids, fibroblasts, and blood-derived macrophages under agitation. Flow cytometry confirmed high expression of eight lectins, with four upregulated in TAMs. Survival analyses associated CLEC4E/Mincle, CLEC6A/Dectin-2, and CD209/DC-SIGN with poorer outcomes. As CLEC4E and CLEC6A converge on FcRγ/SYK signaling, this shared node was selected for pathway-level pharmacological perturbation. R406 reduced SYK phosphorylation and induced transcriptional remodeling in TAMs, with attenuation of immunosuppressive macrophage features, reduced CD204/CD206, increased HLA-DR, and remodeling of the soluble-factor profile. These findings define a myeloid lectin framework associated with BC immunosuppression and identify SYK as a candidate targetable node for remodeling TAMs.

## Introduction

Despite major advances in endocrine and tumor-targeted therapies, breast cancer (BC) still carries a substantial burden of relapse and mortality.[1] Current therapeutic modalities control tumor-intrinsic drivers, but do not address immunosuppressive (IS) tumor microenvironments (TMEs).[2] Moreover, response varies with biological heterogeneity across molecular subtypes and microenvironmental states, which together shape antigenicity, immune infiltration, and checkpoint dependence.[3,4] Immunotherapy has emerged as a complementary strategy to overcome resistance. Yet, most efforts have remained T-cell centric, which limits success outside narrow contexts.[5] These limitations highlight the need for additional axes that reprogram immunosuppression at its source.[6]

Within the myeloid compartment of the TME, tumor-associated macrophages (TAMs) are often abundant and consistently linked to progression and worse prognosis in BC.[7] Functionally, immunosuppressive TAMs adopt immunoregulatory programs with reduced co-stimulatory tone, dampening effector immunity and promoting tissue remodeling and angiogenesis via anti-inflammatory and metabolic cues, thereby contributing to therapeutic resistance.[8]

Despite their promise, TAM-targeting immunotherapies have shown limited efficacy as monotherapies: CD47–SIRPα blockade has shown limited activity in solid tumors;[9] CSF1R or CCL2/CCR2 inhibition, aimed at reducing TAM recruitment, has been hampered by pathway redundancy and compensatory signaling, leading to immune escape.[10,11] Collectively, these observations support a shift from TAM depletion to TAM reprogramming and highlight the need for novel targets that coordinate multiple suppressive pathways, enabling multi-axis suppression to be addressed through combination therapies.[12]

Glycosylation, being the most abundant and structurally diverse post-translational modification, pervasive across cell-surface and secreted proteins and extending to glycolipids and proteoglycans, provides such an avenue.[13] Cancer cells remodel their surface glycome, exhibiting characteristic glycan alterations, such as truncated O-glycans (Tn/STn), hypersialylation, core fucosylation, and MGAT5-driven β1,6-branched N-glycans. These tumor-associated glycans reshape receptor signaling and immune recognition. They are high-affinity ligands for lectin receptors on myeloid cells, creating “glyco-immune checkpoints” that tune macrophage activation.[14] Lectins bind carbohydrates through carbohydrate-recognition domains (CRDs) and act as carbohydrate pattern sensors that can enforce or relieve immune suppression within the TME. [11,15]

In BC, specific glycan–lectin axes on macrophages have been linked to immunoregulation and disease progression: for example, CD24–SIGLEC-10 imposes an anti-phagocytic signal[16]; sialylated, truncated MUC1 (Tn/STn)–SIGLEC-9 skews toward IL-10–rich programs[17]; DC-SIGN engagement by fucosylated tumor glycoproteins elevates VEGF/IL-10 and dampens antigen presentation,[18] and MGL (CLEC10A) recognition of Tn/STn motifs promotes tolerogenic signaling.[19] However, these reports are largely target-centric and cover a narrow subset of lectins.[20] This gap motivates a comprehensive, macrophage-focused assessment across the transmembrane immune lectin repertoire to define druggable nodes that complement existing strategies.

Here, we took a myeloid-focused approach, combining multi-omics prioritization with manual curation, to map the immunosuppressive lectin landscape in BC. Using a clinically annotated, population-scale cohort (SCAN-B)[21], we identified an immunosuppressive tumor cluster with a validated immune classifier.[22] We then integrated bulk differential expression with single-cell myeloid references[23] and upstream-regulator inference to prioritize druggable transmembrane immune lectins. Identified candidates were evaluated at the protein level in a human reconstructed 3D heterotypic TME model that reproducibly generates immunoregulatory TAMs.[24] Finally, based on the convergence of CLEC4E/Mincle and CLEC6A/Dectin-2 on FcRγ/SYK signaling, we used pharmacological SYK inhibition as a pathway-level perturbation. SYK inhibition attenuated the immunosuppressive TAM markers and induced coordinated transcriptional and metabolic remodeling, supporting a functional contribution of this shared downstream node to TAM polarization. Rather than focusing on a single mediator, this framework maps glycan-sensing immune checkpoints on myeloid cells and nominates myeloid lectin and associated signaling pathways as candidate targets for macrophage modulation in BC.

## Results

### Bulk Transcriptomics Reveals Dysregulated Lectins in Immunosuppressive Tumors

Using a published immune index[22] to stratify the SCAN-B cohort (n=3,207), we identified 1,397 Immunosuppressive (IS; 43.6%) and 1,810 Non-IS (56.4%) tumors. An unsupervised PCA (Fig. 1A) on the classifier genes showed clear structure: PC1 (16.8% variance explained) largely separated Non-IS, while PC2 (14.0%) captured the immunosuppressive gradient, yielding a characteristic split on an orthogonal projection, consistent with a continuum of immune states rather than a binary partition.

**Figure 1.**
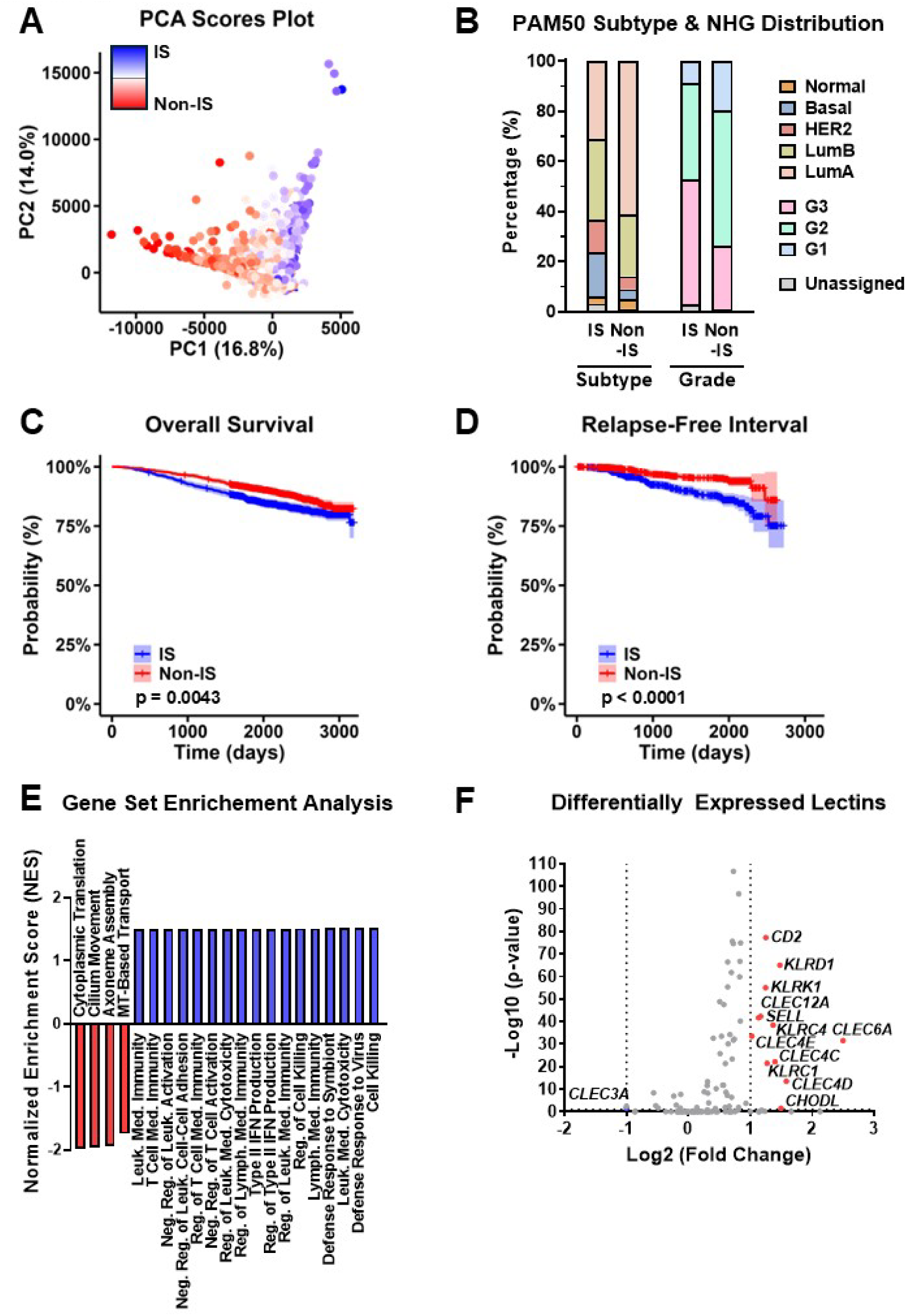
Bulk Transcriptomics reveals dysregulated lectins in immunosuppressive tumors. Principal component analysis (PCA) score plot from the expression of the 652 genes used in immunosuppressive cluster classification for the 3207 samples, colored by gradient scale on index value and centered on threshold value (A). Distribution of samples according to PAM50 subtype and Nottingham histological grade (NHG) between IS and Non-IS clusters (B). Kaplan-Meier overall survival (C) and relapse-free interval (D) analysis curves comparing samples in IS vs Non-IS clusters. Gene set enrichment analysis (GSEA) for Gene Ontology: Biological Processes in IS vs Non-IS clusters (E). Differential gene expression analysis (DGEA) of the 112 lectins in IS vs Non-IS clusters (F).

Clinicopathologic parameters (Fig. 1B) aligned with the discrimination of clusters based on expected immunosuppressive features. The IS cluster was enriched for Basal-like (>3-fold vs Non-IS) and HER2-enriched samples (2.5-fold), whereas Luminal A was 2-fold more frequent in Non-IS. High histologic grade (G3) was 2-fold more common in IS, with G1 and G2 being relatively over-represented in Non-IS (2.3-fold and 1.4-fold, respectively). Other variables (age at diagnosis, nodal status, Ki-67, endocrine therapy, chemotherapy, and histologic subtype) did not differ significantly between immune clusters (Supplementary Fig. S2).

Survival analyses indicated poorer outcomes in the IS cluster, with Kaplan–Meier curves for overall survival (OS, Fig. 1C) and relapse-free interval (RFI, Fig. 1D) consistently lower survival probabilities compared with the Non-IS cluster. Although effect sizes were modest (expected in treated, heterogeneous patient cohorts), the persistent separation and directionality were consistent with a protumor state in the IS cluster.

Further gene-set enrichment analysis (Fig. 1E) highlighted the IS cluster co-enriched immune effector/IFN-γ-response programs, together with immune-regulatory/exhaustion signatures (e.g., negative regulation of leukocyte and T cell activation), consistent with an “immune-rich yet immunosuppressed/exhausted” state. In contrast, Non-IS tumors were enriched for non-immune, cell-intrinsic programs, compatible with immune exclusion and a comparatively “cold” TME. Importantly, regulatory/exhaustion enrichments in IS tumors indicate an immune-inhibitory state rather than stronger activation, compatible with potential myeloid lectin-driven control.

Having established the IS assignment, DGEA (Fig. 1F) was performed, focusing results on the 112 transmembrane lectins identified from a curated set (C-type and I-type). We identified 13 potentially discriminant lectins: 12 upregulated in the IS Cluster, and 1 upregulated in Non-IS, based on bulk tumor profiles, reflecting the composite TME gene expression

### Single-Cell Mapping Shows a Myeloid-High Lectin Signature in Breast Cancer TME

To profile overexpressed myeloid lectins, a scRNA-Seq cohort comprising samples from 88 BC patients[23] was used (Supplementary Fig. S3). The 112 transmembrane lectins identified from the curated set were ranked by expression within the myeloid compartment. The top quartile-ranked lectins were considered overexpressed, having an above-average number of transcripts and clearly separating from the remaining lectins (Fig. 2A). This analysis identified 27 myeloid lectins. Further analysis aimed to understand if their expression associated with specific immunosuppressive myeloid phenotypes.

**Figure 2.**
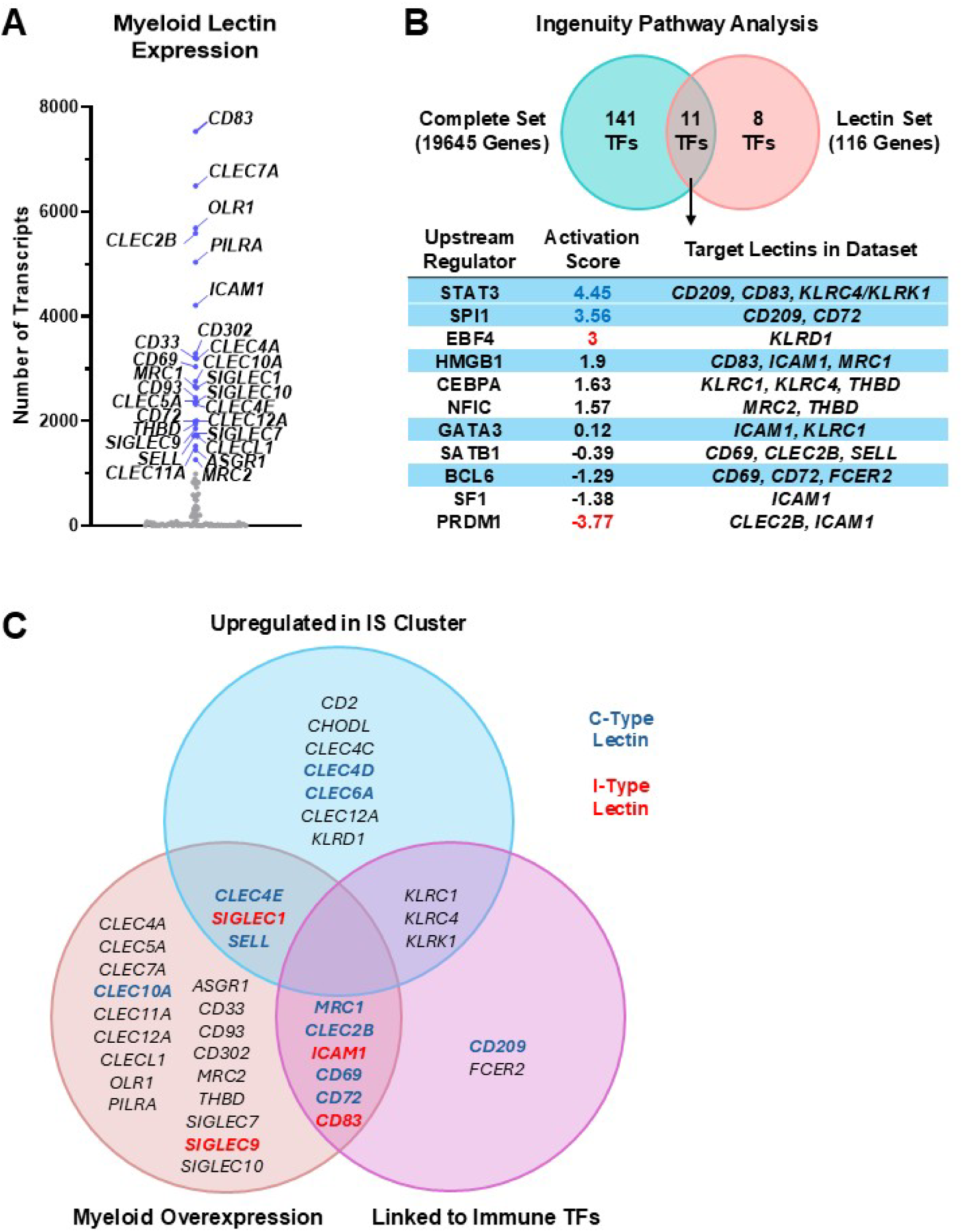
Lectin signature in myeloid cells and immune regulatory programs for orthogonal integration and target selection. Mapping of lectin expression by number of transcripts in myeloid cells, highlighting the top 25% expressing ones in blue (A). Identification of transcription factors (TFs) identified by Ingenuity Pathway Analysis (IPA) for the complete set of genes and the lectin set, and characterization of lectins associated with each transcription factor (B). TFs associated with immune biological processes by Gene Ontology are highlighted in blue. Integration of the three prioritization axes, supplemented by expert-curation(C). Relevant lectins are highlighted in bold, with C-type lectins in blue and I-type lectins in red.

### Regulatory Inference Links Immune Programs to Lectin Transcription

In parallel, we mapped potential immune-related transcriptional drivers of lectin expression. Upstream regulator analysis was performed by Ingenuity Pathway Analysis (IPA, QIAGEN Inc.) on two gene sets from SCAN-B: (i) the complete gene set comprising 19,645 genes, and (ii) a lectin filtered set containing 112 genes. The whole-transcriptome analysis identified 152 significant upstream regulators, while the lectin-specific analysis yielded 19 (Fig. 2B). Intersection of these two sets revealed 11 upstream regulators common to both analyses, indicating a subset of regulators consistently associated with lectin expression but supported by genome-wide context. Amongst these, the strong STAT3 activation supports an immunosuppressive myeloid program consistent with the established role of STAT3 in coordinating tumor-associated immune evasion through macrophages and other immune-cell populations.[25]

Functional annotation of these 11 upstream regulators using the UniProt Knowledgebase biological process terms (GO:BP) identified 5 regulators linked to immune-related processes (highlighted in Fig. 2B). Extraction of lectin genes predicted by IPA to be regulated by these immune-related transcription factors resulted in a final list of 12 lectins potentially under immune-specific transcriptional control.

### Multi-Evidence Prioritization and Curation Define a Lectin Panel for Validation

Building on the previous findings, we combined evidence across three axes to prioritize lectins (i) IS-associated upregulation in bulk, (ii) myeloid-compartment enrichment in single-cell data, and (iii) linkage to immune-related upstream regulators (Fig. 2C). No single lectin satisfied all three axes concurrently. We therefore retained lectins concordant across any two evidence streams, with a specificity guard excluding those lacking myeloid expression. This yielded two intersection sets: IS upregulation ∩ myeloid overexpression and myeloid overexpression ∩ immune regulatory. Within the IS upregulation ∩ myeloid overexpression set, *CLEC4E*/Mincle emerged. We then curated the panel with additions grounded in prior mechanistic and clinical knowledge. Since *CLEC4E* depends on *CLEC4D*/MCL for optimal signaling and *CLEC6A*/Dectin-2 shares the FcRγ–SYK–CARD9 pathway,[26] we added *CLEC4D* and *CLEC6A* to capture cluster-level cooperation. Finally, we included *SIGLEC9*, *CD209*/DC-SIGN, and *CLEC10A*/MGL as canonical myeloid lectins to serve as benchmark comparators and to leverage knowledge from previous reports.[15] Together, this integration yielded a 12-lectin panel for validation at the protein level.

### Lectin Surface Expression in the 3D IS-TME Defines a Panel for Functional Validation

To evaluate hit lectins in a physiologically relevant context, we used a previously established 3D heterotypic TME model (Fig. 3A). Tumor spheroids of BC cell lines were co-cultured with healthy-donor PBMC-derived monocytes and primary human dermal fibroblasts within alginate microcapsules. This biochemically inert, shear-protective matrix supports ECM accumulation and diffusion of oxygen, nutrients, and secreted factors.[24,27] Three cell lines from different BC subtypes were used: HCC1806 (TNBC – Basal-Like), MDA-MB-231 (TNBC – Claudin Low) and BT474 (Luminal B), to ensure any findings are not cell-line-or subtype-dependent.

**Figure 3.**
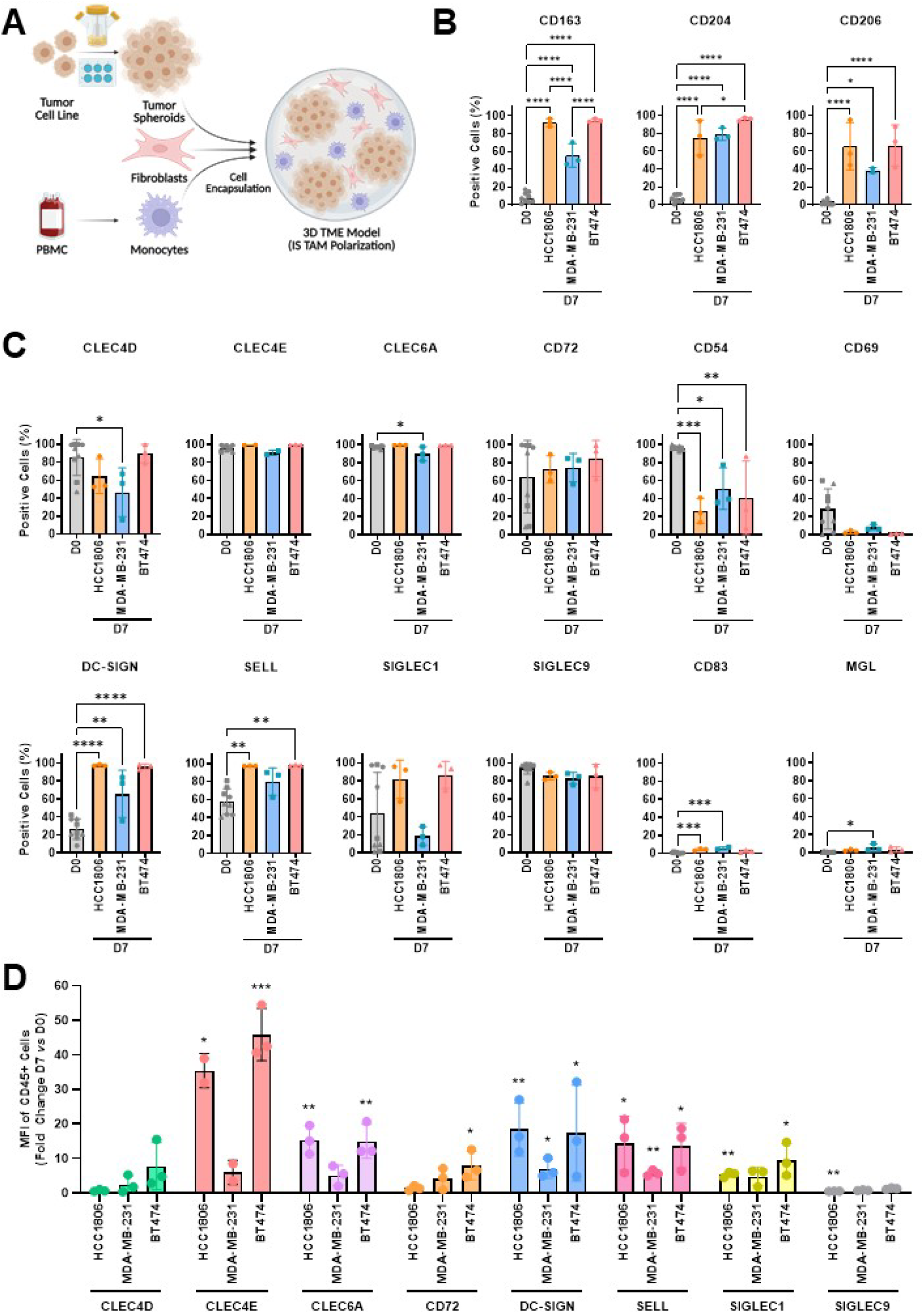
3D IS-TME prioritizes surface-expressed myeloid lectins. Experimental design of the 3D IS-TME model, comprising aggregated tumor spheroids, fibroblasts and PBMC-derived monocytes, encapsulated in an alginate polymer (A). Percentage of positive cells (relative to live or CD45+ cells) for CD163, CD204 and CD206 (B) or selected lectins (C) from monocytes (Day 0) and recovered macrophages (Day 7) in co-culture with HCC1806, MDA-MB-231 and BT474 cells in the 3D IS-TME. Fold-change in the MFI of CD45+ cells at Day 7 *vs* Day 0 for each selected lectin in monocytes/macrophages across 3D IS-TME with three different cell lines (D). Statistical significance was determined using one-way ANOVA with Tukeýs multiple comparisons test (B and C) or one sample ratio t-test (D), with p < 0.05 (*), p < 0.01 (**), p < 0.001 (***) and p < 0.0001 (****). N=3 independent biological donors for each cell line.

By day 7 in co-culture, monocytes differentiated into macrophages, as indicated by increased CD14 and CD64 expression (MFI) relative to day 0 (Supplementary Fig. S4A). Moreover, an anti-inflammatory phenotype was identified by the robust increase in the percentage of CD163^+^/CD204^+^/CD206^+^ cells (Fig. 3B), with immunoregulatory features (Supplementary Fig. S4B-C). With an IS-TAM context established, we assessed surface expression of the 12 prioritized lectins. To identify the most relevant lectin candidates, we first applied a prevalence-based filter (Fig. 3C), retaining lectins that were expressed by a high proportion of cells at day 7, irrespective of changes in expression relative to monocytes (day 0). Eight lectins met this criterion: CLEC4D, CLEC4E, CLEC6A, CD72, and SIGLEC9 by maintaining high detection, and DC-SIGN, SELL, and SIGLEC1 by increasing detection from day 0. In contrast, CD54 declined over time, the CD69 level was low, and CD83 and MGL were undetectable at both time points; thus, these four lectins were excluded. Second, we filtered by induction level (Fig. 3D), examining the D7/D0 MFI fold-change to prioritize lectins whose detection intensity increased in the immunosuppressive context. Indeed, the largest fold-changes at day 7 were observed in BT474 and HCC1806 for CLEC4E (∼35-45-fold), CLEC6A (∼15-fold), DC-SIGN (∼18-fold), and SELL (∼14-fold); increases were more modest in MDA-MB-231, in accordance with its weaker immunosuppressive TAM polarization (Fig. 3B).

Exploratory survival analyses in SCAN-B further showed that higher expression of *CLEC4E*, *CLEC6A,* and *CD209* (DC-SIGN) at the tumor level correlated with shorter overall survival and relapse-free interval, consistent with an adverse immunosuppressive milieu, whereas *SELL* showed the opposite association (Supplementary Fig. S5). Together, these data highlight CLEC4E, CLEC6A, and CD209/DC-SIGN as surface-induced lectins with adverse clinical association in the IS-TME model.

### SYK Inhibition Attenuates Immunosuppressive TAM Polarization

Interestingly CLEC4E and CLEC6A converge on FcRγ/SYK-dependent signaling. To test whether inhibition of this shared downstream node could remodel TAM polarization, we performed a proof-of-concept pathway-level pharmacological challenge with R406, the active metabolite of fostamatinib and known inhibitor of SYK kinase activity.[28]

SYK expression and inducible phosphorylation were confirmed in macrophages, but not in BT474 tumor cells, using HAGG stimulation as a positive control (Supplementary Fig. S6A-B). R406 toxicity profiling defined 0.2 µM as a suitable concentration for downstream experiments (Supplementary Fig. S6C). In BT474–monocyte co-cultures, R406 reduced the pSYK/SYK ratio by 52%, confirming inhibition of SYK phosphorylation (Fig. 4A-B).

**Figure 4.**
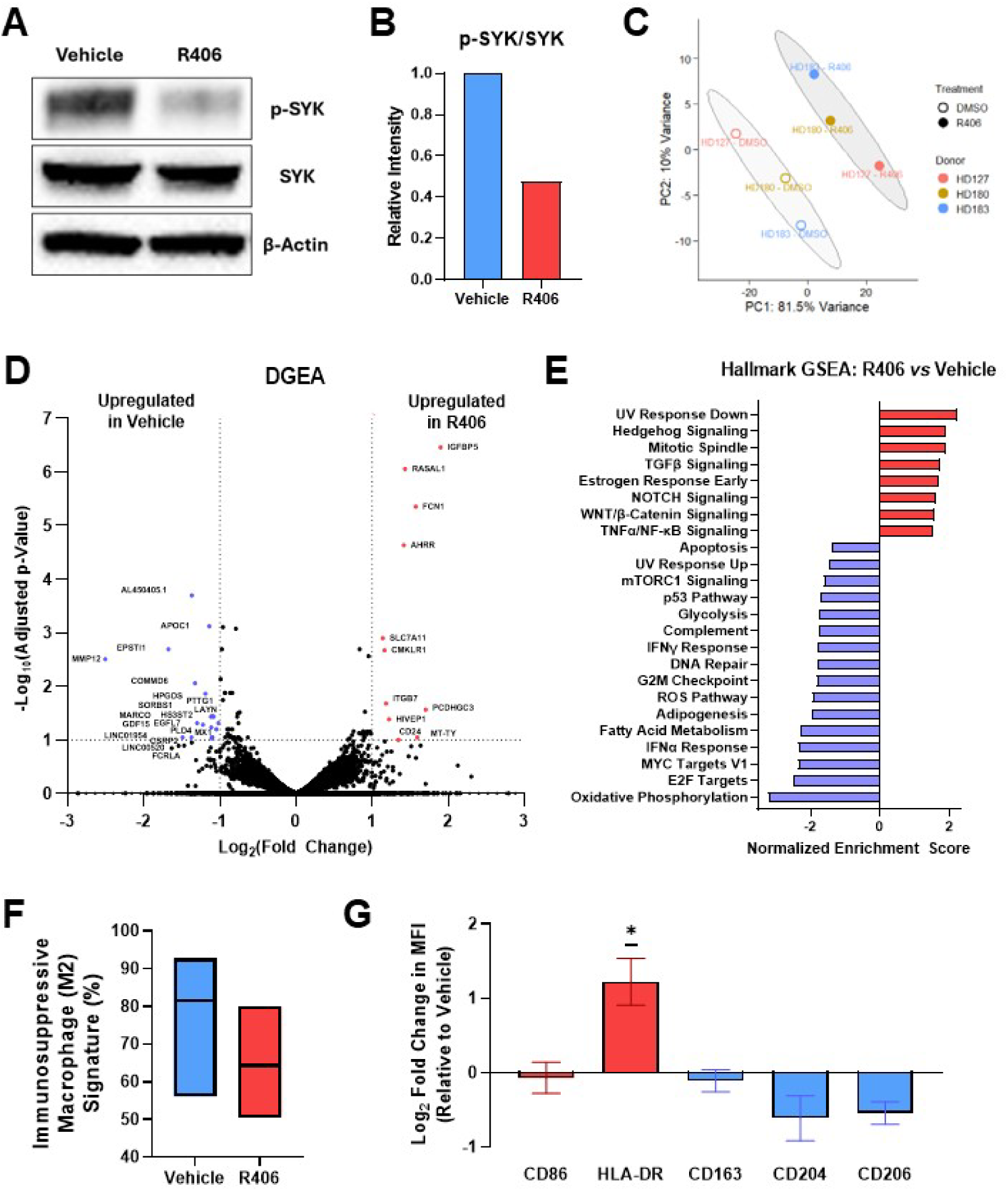
SYK inhibition attenuates immunosuppressive TAM polarization. Western blot analysis of p-SYK, total SYK and β-actin in vehicle-and R406-challenged BT474–monocyte co-cultures (A), with quantification of the p-SYK/SYK ratio (B). Cropped blot; full Length blots are presented in Supplementary Fig. S7. PCA score plot of bulk RNA-Seq from sorted TAMs after donor-effect correction, showing separation of DMSO-and R406-treated samples; shaded ellipses indicate 95% data regions for each treatment group (C). Differential gene expression analysis of sorted TAMs from R406 or vehicle challenge, highlighting genes downregulated (blue) or upregulated (red) upon SYK inhibition (D). Hallmark GSEA of R406-treated TAMs, showing pathways enriched in R406 or reduced by R406 relative to vehicle (E). *quanTIseq* estimation of immunosuppressive macrophage (M2) signature abundance in vehicle-and R406-treated TAMs (F). Flow cytometry validation of TAM polarization markers upon R406 treatment, showing changes in HLA-DR, CD86, CD163, CD204 and CD206 relative to vehicle controls (G). Statistical significance was determined using a one-sample t-test (G), with p < 0.05 (*). N=1 (A-B) or N=3 (C-G) independent biological donors.

TAMs sorted from R406-treated co-cultures were analyzed by bulk RNA-Seq. PCA showed clear treatment-associated separation after donor effect removal, indicating that SYK inhibition induced a reproducible transcriptional shift despite donor-driven variability (Fig. 4C; Supplementary Fig. S6D). Differential expression analysis showed that R406 challenge downregulated several genes associated with immunosuppressive and tumor-promoting TAM phenotypes, including *MMP12*, *MARCO*, *HPGDS*, *GDF15*, *PTTG1*, *LAYN*, and *APOC1* (Fig. 4D). [29–35] Conversely, R406 upregulated *AHRR*, *ITGB7*, *FCN1*, *CMKLR1*, *RASAL1*, and *HIVEP1* (Fig. 4D), genes linked to anti-tumorigenic or less immunosuppressive immune contexts. [36–41]

GSEA further supported a coordinated macrophage remodeling response. R406 reduced oxidative phosphorylation, fatty acid metabolism, glycolysis, ROS, mTORC1, MYC, E2F, interferon, and complement programs, suggesting attenuation of a metabolically active IFN/complement-rich TAM state (Fig. 4E). In parallel, TNFα/NF-κB, TGFβ, NOTCH, and WNT/β-catenin signaling pathways were enriched following R406 challenge, indicating selective inflammatory rewiring.

Consistent with this transcriptional shift, we observed a reduction in the immunosuppressive macrophage signature upon R406 treatment (Fig. 4F), as assessed by *quanTIseq* deconvolution.[42] Flow cytometry validation confirmed a decrease in the immunoregulatory markers CD204 and CD206, together with increased HLA-DR expression, consistent with a shift towards a less immunosuppressive macrophage phenotype (Fig. 4G). Complementary Olink profiling of the co-culture secretome identified reductions in CCL7, MMP-1 and IL-1β, consistent with attenuation of tumor-supportive recruitment, matrix-remodeling, and inflammatory outputs (Supplementary Fig. S6E). Remarkably, these changes were not accompanied by reduced viability in either BT474 tumor cells or macrophage populations, suggesting that the effects of R406 were primarily driven by its pharmacological activity rather than by nonspecific cytotoxicity (Supplementary Fig. S6F). Together, these data indicate that pharmacological inhibition of SYK phosphorylation attenuates selective immunosuppressive TAM features, providing proof-of-concept evidence that targeting this shared signaling node can promote macrophage remodeling in BC.

## Discussion

This study integrates population-scale profiling with cell-type resolution and pathway-level perturbation to examine the impact of myeloid lectins in BC immunosuppression. Using an immune classifier to stratify the SCAN-B cohort, we observed an immunosuppressive spectrum associated with adverse clinicopathologic features and outcomes. Within this context, differential gene expression pointed to a restricted set of transmembrane lectins enriched in immunosuppressive BC. Single-cell analyses localized these lectins predominantly to the myeloid compartment, and upstream-regulator inference highlighted immune-linked transcriptional programs associated with lectin expression. In a reconstructed 3D TME that yields macrophages with immunoregulatory features, surface-level validation prioritized lectins associated with adverse clinical outcome. Finally, pharmacological inhibition of SYK, a shared downstream effector of CLEC4E/Mincle and CLEC6A/Dectin-2, attenuated immunosuppressive TAM polarization. Together, the results suggest that myeloid lectin expression marks immunosuppressive TME states in BC and nominate lectin-associated signaling for further functional investigation.

The SCAN B analysis recapitulates a pattern seen across solid tumors in which effector type programs coexist with regulatory or exhaustion signals and do not necessarily translate into improved outcomes. The enrichment of Basal like and HER2 enriched subtypes within the IS group and the modest but persistent separation of survival curves are consistent with prior population scale studies showing that myeloid dominant inflammation can coincide with poorer prognosis in BC.[22,43,44] These observations provide clinical context for the subsequent prioritization of myeloid expressed lectins.

Our primary analysis centered on the two classes of transmembrane immune lectins. I-type lectins (which include *SIGLEC1*, *SIGLEC9*, *CD83,* and *ICAM1*) recognize sialylated ligands via Ig-like domains, and typically carry immunoreceptor tyrosine-based inhibitory motifs (ITIMs) that recruit SHP-1/2 on engagement, sensing self-associated molecular patterns (SAMPs) and dampening responses to pathogen-associated (PAMPs) and damage-associated molecular patterns (DAMPs).[45–47] In contrast, C-type lectin receptors (CLRs) bind diverse glycans through Ca²-dependent CRDs and are commonly coupled to immunoreceptor tyrosine-based activation motifs (ITAMs) (e.g., via FcRγ), and signal through SYK–CARD9 to tune DAMP-driven inflammation in a context-dependent manner.[46,48]

Combining multi-omics prioritization with expert curation across the transmembrane lectin panel, we identified 12 candidate immunosuppressive lectins for validation. The myeloid-expression filter excluded candidates primarily expressed by NK cells or CD8⁺ T-cell subsets, preserving the macrophage-focused scope of the analysis.[49] The 3D co-culture model offers a pathophysiology-relevant setting in which monocytes differentiate into macrophages exhibiting high CD163/CD204/CD206 with preserved antigen-presentation and limited co stimulation-features characteristic of tolerogenic TAMs.[8,12] Notably, this occurred from endogenous co-culture cues, such as polarizing cytokines, rather than exogenous stimuli. Thus, it implicated cancer-and fibroblast-derived factors as sufficient drivers, consistent with a self-organized IS-TME. Differences in amplitude of polarization across cell lines (stronger immunosuppressive polarization with HCC1806 and BT474 than with MDA-MB-231), likely reflect intrinsic biology; the mesenchymal/EMT-like MDA-MB-231 showed a comparatively weaker immunosuppressive shift.[50]

Among the surface lectins induced in the 3D co-culture model, CLEC4E/Mincle, CLEC6A/Dectin-2, and CD209/DC-SIGN showed the strongest associations with poorer overall survival in the SCAN-B cohort.

CD209/DC-SIGN has been described on alternatively-activated and tumor-associated macrophages and may contribute to an immunosuppressive cytokine environment.[18] By contrast, CLEC4E/Mincle and CLEC6A/Dectin-2 are best characterized as activating FcRγ-coupled pattern-recognition receptors: CLEC4E recognizes microbial and damage-associated ligands, whereas CLEC6A is principally established in antifungal immunity.[51–53] Notably, CLEC4E has also been shown to maintain pro-tumoral TAM activity through SYK–NF-κB signaling, while direct evidence for CLEC6A in TAM biology remains limited.[54]

The convergence of CLEC4E and CLEC6A on FcRγ/SYK-dependent signaling provided the rationale for pathway-level perturbation with R406, allowing us to test whether inhibition of this shared downstream node could attenuate immunosuppressive TAM features.[28]

The R406 challenge supported this hypothesis, but also suggested that SYK inhibition does not simply convert TAMs into a canonical pro-inflammatory state. Instead, R406 reduced an immunosuppressive macrophage signature, decreased CD204/CD206, and increased HLA-DR expression, while transcriptomic analysis suggested broader remodeling of metabolic, interferon/complement, and inflammatory signaling programs. At the transcriptional level, R406 reduced several genes previously linked to tumor-promoting or immunosuppressive myeloid states, including *MARCO*, *APOC1*, *HPGDS*, *GDF15*, *MMP12*, *LAYN*, and *PTTG1*.[29–35] In contrast, genes increased upon R406 challenge, including *AHRR*, *ITGB7*, *FCN1*, *CMKLR1*, *RASAL1*, and *HIVEP1*, point to a distinct macrophage-remodeling program involving inflammatory regulation, immune trafficking, and context-dependent tumor-suppressive pathways.[36–41] Together, these changes support a shift away from selected adverse TAM features, although not a complete macrophage repolarization. At the secretome level, reduction in CCL7, MMP-1 and IL-1β further suggest that SYK inhibition may attenuate myeloid-recruiting, matrix-remodeling, and tumor-promoting inflammatory functions.[55–57]

This distinction is important as CLR-SYK signaling is classically associated with myeloid activation in response to pathogen or tissue-damage signals.[53,58] Still the TME imposes chronic inflammation that can redirect activating pathways toward immunoregulatory phenotypes.[54,59,60] Thus, our data support a context-dependent view of CLEC4E/CLEC6A-SYK signaling in TAMs, where inhibition of a shared downstream node can reduce immunosuppressive TAM features without globally suppressing or uniformly activating macrophages.

Most translational efforts to remodel the myeloid compartment have targeted recruitment and survival (e.g., CCR2/CCR5 or CSF1R)[61,62] or phagocytosis checkpoints (e.g., CD47–SIRPα)[63]. Our findings point to myeloid lectin biology and CLR-associated signaling as a complementary axis, with potential advantages for patient selection and combinatorial pharmacological design. The observation that several lectins are readily detectable on the myeloid cell surface in the context of an immunosuppressive microenvironment, supports their development as immunohistochemical or flow-cytometry-based biomarkers. Clinically, a composite “lectin score” could be explored alongside PD L1 or myeloid gene signatures to identify inflamed but suppressed tumors that may benefit from myeloid directed combinations. Therapeutically, strategies that modulate CLR-associated signaling, lectin engagement, or tumor glycosylation merit investigation, particularly in combination with PD 1/PD L1 blockade. These hypotheses warrant validation in models incorporating spatial organization, tumor glycosylation, and immune readouts, suitable for early phase translational studies.

Some limitations should be considered. Bulk transcriptomics reflect signals from mixed cell populations, and although single-cell analyses supported myeloid localization, the datasets are not patient-matched. The 3D co-culture model lacks adaptive lymphoid compartments, notably T cells. As such, the net effect of lectin modulation on T-cell activation and cytotoxicity remains to be defined. The R406 challenge was designed as a pathway-level perturbation of SYK-dependent signaling rather than as a receptor-specific intervention. Therefore, while the results support the functional relevance of the CLEC4E/CLEC6A–SYK axis, the individual contributions of each CLR to the observed effects could not be delineated within the scope of this study. Furthermore, the ligands and cellular sources mediating lectin engagement in the TME were not investigated and warrant future study. Finally, survival associations were derived from transcript levels in heterogeneous bulk tissue and require further validation at the protein and spatial levels in independent cohorts.

In conclusion, by integrating population level stratification, single cell localization, and pathway-level perturbation, we identify a myeloid lectin program linked to immunosuppressive BC microenvironments. CLEC4E, CLEC6A and CD209/DC-SIGN emerged as surface-induced lectins with adverse clinical associations, while SYK inhibition attenuated selected immunosuppressive TAM feature, highlighting shared CLR-associated signaling as a potential mechanism of macrophage remodeling. These findings further support the development of lectin based biomarkers and provide a rationale for evaluating lectin modulating strategies, alone or in combination with checkpoint blockade, as candidate therapies for myeloid remodeling in BC.

## Materials and Methods

### Bioinformatics & Computational Analysis of Public Breast Cancer Datasets

All analyses were performed in R (v4.2.2) within the RStudio environment (v2025.5.0.496), using base R functions and the *tidyverse* package suite (v2.0.0; including *dplyr* v1.1.4, *tidyr* v1.3.1, *ggplot2* v3.5.1).

### Public Bulk RNA Sequencing Data

Bulk RNA sequencing (RNA-Seq) data from SCAN-B cohort [21], comprising 3,207 clinically annotated BC patient samples, was downloaded from Gene Expression Omnibus (GEO) as a gene-level transcripts *per* million (TPM) expression matrix.

### Immunosuppressive Classification

Immune subtypes were assigned to each sample using a scoring model.[22] Genes from the classifier panel were extracted from the SCAN-B TPM expression matrix, and TPM-normalized expression values were multiplied by the corresponding published LASSO regression coefficients. Weighted expression values were summed to calculate an immune index for each sample. Samples with an index above the threshold (1.206538657) were classified as Immunosuppressive (IS), and those below as Non-Immunosuppressive (Non-IS).

### Principal Component Analysis (PCA)

PCA was performed using mean-centered TPM expression values of the classifier gene panel using the *prcomp* function. PCA scores for the first two principal components were plotted with samples mapped according to a gradient-capped immune index, where limits were set to the 1^st^ and 99^th^ percentiles.

### Differential Lectin Expression Analysis

Differential Gene Expression Analysis (DGEA) was performed using unpaired two-tailed t-tests to compare IS and Non-IS clusters. For each gene, mean expression, log_2_ fold-change (log_2_FC), and Bonferroni-adjusted p-values were calculated. Genes with |log_2_FC|≥1 and adjusted p-value<0.1 were considered significant. Results were filtered using a curated lectin list from the HumanLectome section of the UniLectin database,[64] comprising all confirmed and predicted C-type and I-type lectins (Supplementary File S2).

### Gene Set Enrichment Analysis (GSEA)

Gene Set Enrichment Analysis (GSEA) was performed using *gseGO* function from *clusterProfiler* package (v4.16.6), with a ranked list of all genes ordered by a composite score (log FC×-log (adjusted p-value)). Analyses were conducted against Gene Ontology Biological Processes (GO:BP) using human gene annotations from *org.Hs.eg.db* (v3.20.0). P-values were adjusted for multiple testing using the Benjamini–Hochberg method, and gene sets with an adjusted p-value<0.05 were considered significantly enriched. Results were visualized using the *gseaplot* function from the *enrichplot* package (v1.26.6).

### Ingenuity Pathway Analysis (IPA)

Differentially expressed genes from the bulk RNA-Seq analysis (symbol, fold-change, and p-value) were uploaded into Ingenuity Pathway Analysis (IPA, Qiagen). Genes with p-values>0.05 were excluded. Upstream regulator predictions were filtered for transcription factors and nuclear receptors. Significance was assessed using the p-value of overlap (<0.0001) and activation z-score (|z|>4).

### Single-Cell RNA Sequencing Analysis

An integrated single-cell RNA sequencing (scRNA-Seq) atlas of the primary BC TME comprising 236,363 cells from 119 patient biopsy samples of 88 patients across eight datasets[23], was downloaded from the Broad Institute Single Cell Portal, together with the corresponding cell-type annotations. Raw count matrices and metadata were imported into *Seurat* package (v5.2.1), and the dataset was reprocessed by normalization, identification of variable features, data scaling, principal component analysis, and UMAP dimensionality reduction, following the workflow described in the original study. Cells annotated as macrophages, monocytes, myeloid-derived suppressor cells (MDSCs), dendritic cells, neutrophils, and mast cells were subsetted to define the myeloid compartment and pooled into pseudo-bulk RNA expression data.

### Survival Analysis

Kaplan–Meier analyses of SCAN-B cohort were performed for overall survival (OS) and relapse-free interval (RFI) using the *survival* and *survminer* packages. Patients were compared according to IS or Non-IS classification or, for single-gene analyses, stratified into high-and low-expression groups using an optimal cutoff determined by maximally selected rank statistics. Survival curves were fitted using *survfit* and compared using two-sided log-rank tests. For single-gene analyses, p-values were adjusted for multiple cutoff testing using the Bonferroni method.

### Cell Culture

BT474 (ATCC; RRID:CVCL_0179), HCC1806 (ATCC; RRID:CVCL_1258), and MDA-MB-231 (ATCC; RRID:CVCL_0062) BC cells (female origin) were cultured in RPMI1640 medium (Gibco), 10% (v/v) FBS (Gibco). Cells were sub-cultured twice a week at 25,000 cell/cm^2^ (BT474), 15,000 cell/cm^2^ (HCC1806), or 20,000 cell/cm^2^ (MDA-MB-231). Human dermal fibroblasts (Innoprot) were cultured in IMDM (Gibco), 10% (v/v) FBS. Fibroblasts were sub-cultured every 2 weeks, at 5,000 cell/cm^2^. The medium was replaced every week. After thawing, cells were sub-cultured at least twice before aggregation. Both fibloblasts and cancer cell line cultures were maintained in a humidified incubator at 37°C and 5% CO_2_. All cell lines were routinely tested for mycoplasma contamination.

### Cancer Cell Aggregation

BT474 cells were inoculated at 0.2×10^6^ cell/mL in 125 mL spinner vessels (Corning), on magnetic stirring plates with 80 rpm agitation, and aggregated for 1 day, according to our previously established protocols. [65,66] For MDA-MB-231 and HCC1806, cells were seeded at 3×10^6^ cell/well in AggreWell™ 400 plates (StemCell Technologies), following the manufacturer’s instructions, and aggregated for 3 days. Spheroids were recovered from spinner vessels and AggreWell™ plates for subsequent applications.

### Human Peripheral Blood-Derived Monocyte Isolation

Anonymized human buffy coats from healthy donors who had provided written informed consent for the use of their donated blood for research purposes were obtained from the Portuguese Institute for Blood and Transplantation. Isolation of peripheral blood-derived mononuclear cell (PBMCs) from buffy coats and then monocytes from PBMCs was performed as reported previously.[24,27] Briefly, buffy coats were diluted 1:1 in DPBS supplemented with 2% (v/v) FBS and 2 mм EDTA (Promega). Diluted buffy coats were processed by density centrifugation with Lymphoprep™ on SepMate™-50 (StemCell Technologies), and the PBMCs fraction was recovered. Subsequent washing steps and incubation with ACK Lysis Buffer (Gibco) were performed to minimize contamination with platelets and erythrocytes. PBMCs were cryopreserved in liquid nitrogen until use. For each culture, PBMCs were thawed, and monocytes isolated through immunomagnetic negative selection, using the EasySep Human Monocyte Isolation Kit (StemCell Technologies), according to manufacturer’s instructions. Monocytes were used immediately for downstream culture experiments.

### Monocyte-Derived Macrophage Monocultures

For macrophage monocultures, freshly isolated monocytes were seeded at 3×10 cell/cm² in RPMI, 10% FBS, 1% penicillin/streptomycin (P/S), and 50 ng/mL M-CSF (PeproTech). Cells were cultured for 7 days at 37°C and 5% CO, with half-medium replacement at day 3. These monocultures were used for R406 toxicity screening and heat-aggregated IgG (HAGG)-induced SYK activation assays (Supplementary Methods).

### Alginate Microencapsulation and 3D IS-TME Cultures

Alginate microencapsulation of BC multicellular spheroids was performed using an electrostatic encapsulator (VarV1, Nisco) as previously described.[24,65,66] Briefly, spheroids were collected from spinner vessels or AggreWell™ plates, and combined with fibroblasts and monocytes in 1.1% (w/v) Pronova® UP MVG sodium alginate (NovaMatrix), 0.9% (w/v) NaCl. Creast cancer cells, fibroblasts, and monocytes were used at a 1:1:1 ratio, corresponding to 24×10 cells/mL alginate. Alginate suspension was infused at 10 mL/h through a cell encapsulator (NISCO) with a 500 μm nozzle, charged at 5.3 kV, and collected into a 20 mM BaCl_2_, 115 mM NaCl solution. The 3D IS-TME cultures were maintained for 7 days in shake flasks, in RPMI, 10% FBS, 1% P/S, in a Multitron incubation shaker (INFORS HT) under 80 rpm continuous orbital agitation, at 37°C, 5% CO_2_. Half the medium was replaced at day 3. Independent 3D IS-TME cultures were established employing monocytes isolated from three different donors for each cell line.

### BT474–Monocyte Spheroid Co-Cultures and R406 Challenge

For pharmacological SYK inhibition experiments, BT474-monocyte co-cultures were established in round-bottom ultra-low attachment 96-well plates. BT474 cells and monocytes were mixed at a 1:1 ratio and seeded at 5×10 total cells per well. Plates were centrifuged at 300×g for 5 min to induce cell aggregation and spheroid formation. R406 besylate (InvivoGen) was added at cell seeding at a final concentration of 0.2 µM. Vehicle-control cultures received matched DMSO concentration (0.314%, v/v). Co-cultures were maintained for 7 days at 37°C and 5% CO, with half-medium change with fresh R406 and DMSO at day 3. Independent cultures were performed using monocytes from three different donors.

### R406 Toxicity Assessment

R406 toxicity was assessed in monocyte-derived macrophage monocultures. R406 was tested in a 12-point two-fold dilution series ranging from 10 to 0.004 µM, with compound refreshed during the half-medium change. Metabolic activity was assessed by resazurin reduction using PrestoBlue (Invitrogen) after 90 min incubation. IC50 values were estimated in GraphPad Prism (v10.5.0) using a nonlinear four-parameter logistic regression with variable slope.

### Western Blot

Cell lysis and protein extraction were performed in RIPA buffer (Merck) supplemented with cOmplete protease inhibitor cocktail (Sigma-Aldrich) and PhosSTOP phosphatase inhibitor cocktail (Roche). Protein concentration was determined by BCA assay (Invitrogen). Lysates were prepared in reducing sample buffer, denatured at 70°C for 10 min, and 15 µg of protein was loaded per lane. Proteins were separated in NuPAGE 4–12% Bis-Tris gels using MOPS running buffer and transferred to 0.45 µm nitrocellulose membranes. Membranes were blocked 1h at RT in 5% non-fat dry milk in TBST and incubated overnight at 4°C with primary antibodies diluted in 5% BSA/TBST: phospho-SYK Y525/526 (Cell Signaling Technology, #2710; RRID:AB_2197222; 1:1000), SYK (Cell Signaling Technology, #80460T; 1:1000; RRID:AB_2799953), and β-actin (Sigma-Aldrich, #A5441; RRID:AB_476744; 1:5000). HRP-conjugated anti-mouse (Cytiva, #NA931; RRID:AB_772210; 1:500) and anti-rabbit (Cell Signaling Technology, #7074; RRID:AB_2099233; 1:500) secondary antibodies were incubated for 1 h at RT. Detection was performed using Amersham ECL Pro and imaged on an iBright FL1500 system (Invitrogen), with p-SYK and SYK detected on separate membranes prepared from the same samples and β-actin as loading control. Densitometry was performed in FIJI, with p-SYK normalized to total SYK.

### Macrophage Phenotype Analysis by Flow Cytometry

For 3D IS-TME cultures, alginate microcapsules were chelated with 100 mM EDTA in 10 mM HEPES for 7 min under agitation and centrifuged at 50×g, for 2 min to separate single cells from the spheroid fraction.[24] The single-cell suspension, comprising mainly fibroblasts and macrophages, was collected for downstream analysis. For R406-treated co-cultures, spheroids were dissociated with TrypLE Express for 15 min at 37°C under agitation. Single-cell suspensions were incubated sequentially with TruStain FcX Block (BioLegend, #422302) for 10 min, conjugated antibodies (Supplementary Table S1) for 1h, and Ghost Dye Violet 510 (Cell Signaling Technology, #16759863S) for 15 min. Incubations were performed on ice with 2% FBS in DPBS containing Ca^²⁺^ and Mg^²⁺^. Data were acquired at iBET Flow Cytometry Unit in a FACSCelesta Cell Analyzer (BD) for 3D IS-TME experiments or an ID7000 spectral flow cytometer (Sony) for ULA/R406 experiments. Results were analyzed in FlowJo (v10) with marker expression reported as FMO-subtracted MFI. The gating strategy is shown in Supplementary Fig. S1.

### TAM Sorting and RNA Extraction

For TAM sorting, BT474–monocyte ULA co-cultures were dissociated and stained as described for flow cytometry. Live singlet CD45⁺ cells were sorted at iBET Flow Cytometry Unit using CD45-BV605 and Ghost Dye Violet 510 on a Sony MA900 cell sorter with a 100 µm sorting chip in semi-purity mode. Sorted cells were collected directly into RLT buffer (Qiagen) supplemented with β-mercaptoethanol (Sigma), snap-frozen in liquid nitrogen, and stored at −80°C. RNA was extracted using the RNeasy Mini Kit (Qiagen, #74106) with on-column DNase digestion for 15 min. RNA concentration was assessed by NanoDrop, and RNA integrity evaluated using Agilent RNA ScreenTape on a TapeStation 4200 system. Samples with RINe≥8 were used for bulk RNA sequencing.

### Bulk RNA Sequencing of Sorted TAMs

Library preparation, sequencing, and gene-level quantification were performed by Novogene. Briefly, mRNA was enriched using poly-T oligo-attached magnetic beads, libraries were prepared following fragmentation and cDNA synthesis, and sequencing was performed on an Illumina platform. Raw reads were processed with *fastp*, aligned to the reference genome using *HISAT2*, and gene-level counts were generated with *featureCounts*. Raw count matrices were used for downstream analysis.

### Analysis of R406-Challenged TAM RNA Sequencing Data

Raw gene counts were analyzed in R using DESeq2 with a paired design including donor and treatment condition *(∼ donor + condition*), comparing R406 against vehicle. Genes with counts ≥10 in at least three samples were retained, and Wald test p-values were adjusted using the Benjamini–Hochberg method. Adjusted p-value<0.1 was used as exploratory threshold for differential expression. Variance-stabilized counts were used for PCA. For visualization of treatment-associated effects, donor variation was removed using *limma::removeBatchEffect*, while preserving the treatment design. GSEA was performed with *clusterProfiler* using Hallmark gene sets from *msigdbr*, with genes ranked by the DESeq2 Wald statistic. Gene sets with Benjamini–Hochberg adjusted p<0.05 were considered significant. For immune cell-fraction estimation, raw counts were converted to TPM and analyzed using the *quanTIseq* deconvolution package with the TIL10 signature matrix and mRNA scaling enabled.[42]

### Cytokine Secretion

The concentration of 15 soluble factors in the supernatant of vehicle-and R406-treated co-cultures was determined by iBET Analytical Services Unit using O-link® proximity extension assay technology (Thermo Fisher Scientific) with a customized Olink® Flex panel. Briefly, 1 µL of each sample was incubated overnight with antibody pairs carrying unique DNA barcodes. DNA tags were then pre-amplified and quantified by real-time qPCR on the Olink® Signature Q100 system. Results were reported as log2 fold change relative to vehicle condition, normalized by median of each sample.

## Statistical Analysis

Statistical analyses were performed using GraphPad Prism software (v10.5.0), unless otherwise stated. Analyses were performed using at least three independent biological replicates, corresponding to cultures established with monocytes from different PBMC donors. One-way ANOVA with Tukey’s multiple comparisons test was used for comparisons involving more than two groups. Flow cytometry fold-change data were analyzed either as log FC relative to control and tested against zero using a one-sample t-test, or as ratio-normalized values tested against one using a one-sample ratio t-test, as appropriate. Unless otherwise stated, p-value<0.05 was considered statistically significant.

### Ethics Approval and Consent to Participate

The use of anonymized human buffy coats of healthy blood donors from the Portuguese Institute for Blood and Transplantation (Instituto Português do Sangue e da Transplantação, IPST, Lisbon, Portugal) was approved by the ethical review board of IHMT – ITQB – NSL – IGC for the project “Descodificar interações glícido-lectina típicas de macrófagos imunossupressores em Cancro - GLYCO-TAM” (reference “Parecer 16.23”). Written informed consent for blood donation and the research use of the donated material was obtained from all donors by the IPST before sample collection. Buffy coats were provided to the investigators in anonymized form, and the investigators had no access to donor-identifying information. All procedures involving human biological material were conducted in accordance with the principles of the Declaration of Helsinki.

The breast cancer transcriptomic datasets analyzed in this study were publicly available and de-identified. Ethical approval and informed consent were obtained by the investigators responsible for the original studies, as reported in the corresponding publications. The present study involved only secondary analysis of publicly available, de-identified data; therefore, no additional ethical approval or informed consent was required for this analysis.

## Acknowledgments

The authors acknowledge Dr. Olivia Rodrigues and the iBET Flow Cytometry Unit for technical expertise, support with TAM single-cell sorting, and access to state-of-the-art instrumentation, as well as the iBET Analytical Services Unit for Olink-based cytokine profiling and associated technical support. The authors further acknowledge Novogene for providing the RNA sequencing services and initial data processing. We are also grateful to Dr. José Escandell (iBET & ITQB NOVA) and Dr. Nuno Raimundo (Penn State College of Medicine & MIA-UC) for valuable scientific discussions that contributed to this work.

The authors acknowledge the use of ChatGPT (OpenAI, GPT-5.5 Thinking) solely for troubleshooting of coding errors during data analysis, and copyediting and language refinement during manuscript preparation. The study design, experiments, analyses, interpretation, and manuscript content were developed by the authors.

## Authors’ Contributions

Conceptualization: GT, GD, II, CB.

Methodology: GT, GD, SB, CB.

Formal analysis: GT, MP.

Investigation: GT, MP, VC, GD.

Resources: ND, CB.

Visualization: GT, MP.

Funding acquisition: CB.

Supervision: GD, AP, II, CB.

Writing – original draft: GT, CB.

Writing – review & editing: GT, GD, MP, VC, SB, ND, AP, II, CB.

All authors read and approved the final manuscript.

## Data Availability

Public SCAN-B bulk RNA-Seq data from Dalal el at. 2022 (https://doi.org/10.1038/s41598-022-08210-3) are available at the Gene Expression Omnibus (GEO), with access number GSE202203. The immune classifier used to stratify tumors into immunosuppressive and non-immunosuppressive groups is available from Tekpli et al. 2019 (https://doi.org/10.1038/s41467-019-13329-5). The public single-cell breast cancer atlas from Xu et al. 2024 (https://doi.org/10.1016/j.xcrm.2024.101511) used for myeloid lectin mapping is available from the Broad Institute Single Cell Portal, study SCP1039. Bulk RNA-Seq data generated from sorted R406-and DMSO-treated TAMs, namely raw sequencing files and count matrices, is available at GEO under accession number GSE342455. Additional data generated in this study are available from the corresponding author upon reasonable request.

## References

1. Bray, F. et al. Global cancer statistics 2022: GLOBOCAN estimates of incidence and mortality worldwide for 36 cancers in 185 countries. CA. Cancer J. Clin. 74, 229–263 (2024).

2. Allegrezza, M. J. & Conejo-Garcia, J. R. Targeted Therapy and Immunosuppression in the Tumor Microenvironment. Trends Cancer 3, 19–27 (2017).

3. de Visser, K. E. & Joyce, J. A. The evolving tumor microenvironment: From cancer initiation to metastatic outgrowth. Cancer Cell 41, 374–403 (2023).

4. Turner, K. M., Yeo, S. K., Holm, T. M., Shaughnessy, E. & Guan, J.-L. Heterogeneity within molecular subtypes of breast cancer. Am. J. Physiol. - Cell Physiol. 321, C343–C354 (2021).

5. Heater, N. K., Warrior, S. & Lu, J. Current and future immunotherapy for breast cancer. J. Hematol. Oncol.J Hematol Oncol 17, 131 (2024).

6. Retecki, K., Seweryn, M., Graczyk-Jarzynka, A. & Bajor, M. The Immune Landscape of Breast Cancer: Strategies for Overcoming Immunotherapy Resistance. Cancers 13, 6012 (2021).

7. Basak, U. et al. Tumor-associated macrophages: an effective player of the tumor microenvironment. Front. Immunol. 14, 1295257 (2023).

8. Kzhyshkowska, J., Shen, J. & Larionova, I. Targeting of TAMs: can we be more clever than cancer cells? Cell. Mol. Immunol. 21, 1376–1409 (2024).

9. Bouwstra, R., van Meerten, T. & Bremer, E. CD47-SIRPα blocking-based immunotherapy: Current and prospective therapeutic strategies. Clin. Transl. Med. 12, e943 (2022).

10. Tufail, M., Jiang, C.-H. & Li, N. Immune evasion in cancer: mechanisms and cutting-edge therapeutic approaches. Signal Transduct. Target. Ther. 10, 227 (2025).

11. Mantuano, N. R., Natoli, M., Zippelius, A. & Läubli, H. Tumor-associated carbohydrates and immunomodulatory lectins as targets for cancer immunotherapy. J. Immunother. Cancer 8, e001222 (2020).

12. Larionova, I., Kazakova, E., Patysheva, M. & Kzhyshkowska, J. Transcriptional, Epigenetic and Metabolic Programming of Tumor-Associated Macrophages. Cancers 12, 1411 (2020).

13. Zhu, Q., Chen, X., Duan, X., Sun, J. & Yi, W. Targeting glycosylation to enhance tumor immunotherapy. Trends Pharmacol. Sci. 46, 863–876 (2025).

14. Pinho, S. S., Macauley, M. S. & Läubli, H. Tumor glyco-immunology, glyco-immune checkpoints and immunotherapy. J. Immunother. Cancer 13, e012391 (2025).

15. Lopes, N., Correia, V. G., Palma, A. S. & Brito, C. Cracking the Breast Cancer Glyco-Code through Glycan-Lectin Interactions: Targeting Immunosuppressive Macrophages. Int. J. Mol. Sci. 22, 1972 (2021).

16. Barkal, A. A. et al. CD24 signalling through macrophage Siglec-10 is a target for cancer immunotherapy. Nature 572, 392–396 (2019).

17. Rodriguez, E. et al. Sialic acids in pancreatic cancer cells drive tumour-associated macrophage differentiation via the Siglec receptors Siglec-7 and Siglec-9. Nat. Commun. 12, 1270 (2021).

18. Domínguez-Soto, A. et al. Dendritic cell-specific ICAM-3-grabbing nonintegrin expression on M2-polarized and tumor-associated macrophages is macrophage-CSF dependent and enhanced by tumor-derived IL-6 and IL-10. J. Immunol. 186, 2192–2200 (2011).

19. Szczykutowicz, J. Ligand Recognition by the Macrophage Galactose-Type C-Type Lectin: Self or Non-Self?—A Way to Trick the Host’s Immune System. Int. J. Mol. Sci. 24, 17078 (2023).

20. Trindade, G., Palma, A. S. & Brito, C. Transmembrane Lectins in Cancer Immunity: Emerging Drivers of Myeloid-Mediated Immunosuppression. Front. Immunol. 17, (2026).

21. Dalal, H. et al. Clinical associations of ESR2 (estrogen receptor beta) expression across thousands of primary breast tumors. Sci. Rep. 12, 4696 (2022).

22. Tekpli, X. et al. An independent poor-prognosis subtype of breast cancer defined by a distinct tumor immune microenvironment. Nat. Commun. 10, 5499 (2019).

23. Xu, L. et al. A comprehensive single-cell breast tumor atlas defines epithelial and immune heterogeneity and interactions predicting anti-PD-1 therapy response. Cell Rep. Med. 5, 101511 (2024).

24. Domenici, G. et al. Assessing Novel Antibody-Based Therapies in Reconstructive 3D Cell Models of the Tumor Microenvironment. *Adv*. Biol. 8, e2400431 (2024).

25. Yu, H., Kortylewski, M. & Pardoll, D. Crosstalk between cancer and immune cells: role of STAT3 in the tumour microenvironment. Nat. Rev. Immunol. 7, 41–51 (2007).

26. Brown, G. D. & Crocker, P. R. Lectin Receptors Expressed on Myeloid Cells. Microbiol. Spectr. 4, 4.5.09 (2016).

27. Rebelo, S. P. et al. 3D-3-culture: A tool to unveil macrophage plasticity in the tumour microenvironment. Biomaterials 163, 185–197 (2018).

28. Rohila, D. et al. Syk Inhibition Reprograms Tumor-Associated Macrophages and Overcomes Gemcitabine-Induced Immunosuppression in Pancreatic Ductal Adenocarcinoma. Cancer Res. 83, 2675–2689 (2023).

29. Shi, B. et al. The Scavenger Receptor MARCO Expressed by Tumor-Associated Macrophages Are Highly Associated With Poor Pancreatic Cancer Prognosis. Front. Oncol. 11, (2021).

30. Ren, L. et al. Systematic pan-cancer analysis identifies APOC1 as an immunological biomarker which regulates macrophage polarization and promotes tumor metastasis. Pharmacol. Res. 183, 106376 (2022).

31. Trotta, R. et al. Activated T Cells Break Tumor Immunosuppression by Macrophage Reeducation. Cancer Discov. 15, 1410–1436 (2025).

32. Haake, M. et al. Tumor-derived GDF-15 blocks LFA-1 dependent T cell recruitment and suppresses responses to anti-PD-1 treatment. Nat. Commun. 14, 4253 (2023).

33. Kerzeli, I. K. et al. Elevated levels of MMP12 sourced from macrophages are associated with poor prognosis in urothelial bladder cancer. BMC Cancer 23, 605 (2023).

34. Pan, J. et al. LAYN Is a Prognostic Biomarker and Correlated With Immune Infiltrates in Gastric and Colon Cancers. Front. Immunol. 10, (2019).

35. Wei, H., Ma, Y., Chen, S., Zou, C. & Wang, L. Multi-omics analysis identifies PTTG1 as a prognostic biomarker associated with immunotherapy and chemotherapy resistance. BMC Cancer 24, 1315 (2024).

36. Vogel, C. F. A. et al. Targeting the Aryl Hydrocarbon Receptor Signaling Pathway in Breast Cancer Development. Front. Immunol. 12, 625346 (2021).

37. Zhu, Y. et al. ITGB7 Remodels Inflammation and Immune Microenvironment and Enhances Checkpoint Inhibitor-Based Immunotherapy in Pancreatic Cancer. J. Inflamm. Res. 18, 17633–17649 (2025).

38. Zhong, F. et al. Multi-omics evaluation of the prognostic value and immune signature of FCN1 in pan-cancer and its relationship with proliferation and apoptosis in acute myeloid leukemia. Front. Genet. 15, (2024).

39. Li, J.-J. et al. Chemerin suppresses hepatocellular carcinoma metastasis through CMKLR1-PTEN-Akt axis. Br. J. Cancer 118, 1337–1348 (2018).

40. Liu, D., Yang, C., Bojdani, E., Murugan, A. K. & Xing, M. Identification of RASAL1 as a Major Tumor Suppressor Gene in Thyroid Cancer. J. Natl. Cancer Inst. 105, 1617–1627 (2013).

41. Matsumoto, H. et al. HIVEP1 Is a Negative Regulator of NF-κB That Inhibits Systemic Inflammation in Sepsis. Front. Immunol. 12, 744358 (2021).

42. Finotello, F. et al. Molecular and pharmacological modulators of the tumor immune contexture revealed by deconvolution of RNA-seq data. Genome Med. 11, 34 (2019).

43. Jääskeläinen, M. M. et al. High Numbers of CD163+ Tumor-Associated Macrophages Predict Poor Prognosis in HER2+ Breast Cancer. Cancers 16, 634 (2024).

44. Yang, M. et al. Stromal Infiltration of Tumor-Associated Macrophages Conferring Poor Prognosis of Patients with Basal-Like Breast Carcinoma. J. Cancer 9, 2308– 2316 (2018).

45. Raposo, C. D., Canelas, A. B. & Barros, M. T. Human Lectins, Their Carbohydrate Affinities and Where to Find Them. Biomolecules 11, 188 (2021).

46. Rabinovich, G. A. & Croci, D. O. Regulatory Circuits Mediated by Lectin-Glycan Interactions in Autoimmunity and Cancer. Immunity 36, 322–335 (2012).

47. Crocker, P. R., Paulson, J. C. & Varki, A. Siglecs and their roles in the immune system. Nat. Rev. Immunol. 7, 255–266 (2007).

48. Zelensky, A. N. & Gready, J. E. The C-type lectin-like domain superfamily. FEBS J. 272, 6179–6217 (2005).

49. Borrego, F., Masilamani, M., Marusina, A. I., Tang, X. & Coligan, J. E. The CD94/NKG2 family of receptors. Immunol. Res. 35, 263–277 (2006).

50. Suárez-Arriaga, M. C., Méndez-Tenorio, A., Pérez-Koldenkova, V. & Fuentes-Pananá, E. M. Claudin-Low Breast Cancer Inflammatory Signatures Support Polarization of M1-Like Macrophages with Protumoral Activity. Cancers 13, 2248 (2021).

51. Robinson, M. J. et al. Dectin-2 is a Syk-coupled pattern recognition receptor crucial for Th17 responses to fungal infection. J. Exp. Med. 206, 2037–2051 (2009).

52. Yamasaki, S. et al. Mincle is an ITAM-coupled activating receptor that senses damaged cells. Nat. Immunol. 9, 1179–1188 (2008).

53. Geijtenbeek, T. B. H. & Gringhuis, S. I. Signalling through C-type lectin receptors: shaping immune responses. Nat. Rev. Immunol. 9, 465–479 (2009).

54. Li, C. et al. The Mincle/Syk/NF-κB Signaling Circuit Is Essential for Maintaining the Protumoral Activities of Tumor-Associated Macrophages. Cancer Immunol. Res. 8, 1004–1017 (2020).

55. Wang, Q.-M., Lv, L., Tang, Y., Zhang, L. & Wang, L.-F. MMP-1 is overexpressed in triple-negative breast cancer tissues and the knockdown of MMP-1 expression inhibits tumor cell malignant behaviors in vitro. Oncol. Lett. 17, 1732–1740 (2019).

56. Nguyen, H. T. et al. Patient-specific vascularized tumor model: Blocking monocyte recruitment with multispecific antibodies targeting CCR2 and CSF-1R. Biomaterials 312, 122731 (2025).

57. Kaplanov, I. et al. Blocking IL-1β reverses the immunosuppression in mouse breast cancer and synergizes with anti-PD-1 for tumor abrogation. Proc. Natl. Acad. Sci. U. S. A. 116, 1361–1369 (2019).

58. Strasser, D. et al. Syk Kinase-Coupled C-type Lectin Receptors Engage Protein Kinase C-δ to Elicit Card9 Adaptor-Mediated Innate Immunity. Immunity 36, 32–42 (2012).

59. Grivennikov, S. I., Greten, F. R. & Karin, M. Immunity, Inflammation, and Cancer. Cell 140, 883–899 (2010).

60. Mantovani, A., Allavena, P., Sica, A. & Balkwill, F. Cancer-related inflammation. Nature 454, 436–444 (2008).

61. Pant, A. et al. CCR2 and CCR5 co-inhibition modulates immunosuppressive myeloid milieu in glioma and synergizes with anti-PD-1 therapy. Oncoimmunology 13, 2338965 (2024).

62. Duran, C. L. et al. Targeting CSF-1 signaling between tumor cells and macrophages at TMEM doorways inhibits breast cancer dissemination. Oncogene 44, 3297–3309 (2025).

63. Zhang, W. et al. Advances in Anti-Tumor Treatments Targeting the CD47/SIRPα Axis. Front. Immunol. 11, 18 (2020).

64. Schnider, B. et al. HumanLectome, an update of UniLectin for the annotation and prediction of human lectins. Nucleic Acids Res. 52, D1683–D1693 (2024).

65. Estrada, M. F. et al. Modelling the tumour microenvironment in long-term microencapsulated 3D co-cultures recapitulates phenotypic features of disease progression. Biomaterials 78, 50–61 (2016).

66. Santo, V. E. et al. Adaptable stirred-tank culture strategies for large scale production of multicellular spheroid-based tumor cell models. J. Biotechnol. 221, 118–129 (2016).

