## Supplementary File S1 for "Myeloid Lectin Profiling Identifies SYK as a Targetable Signaling Node for Remodeling Immunosuppressive Tumor-Associated Macrophages in Breast Cancer"

### Targeting Myeloid Lectins to Reprogram Immunosuppressive Tumor-Associated Macrophages in Breast Cancer

Gonçalo Trindade [1,2]

Giacomo Domenici [1,2]

Miguel Pinto [1,2]

Viviana Correia [1,2]

Sofia Batalha [1,2]

Nádia Duarte [1,2]

Angelina de Sá Palma [6,7]

Inês Azevedo Isidro [1,2]

Catarina Brito [1,2]

[1] iBET, Instituto de Biologia Experimental e Tecnológica, Oeiras, Portugal

[2] Instituto de Tecnologia Química e Biológica António Xavier, Universidade NOVA de Lisboa, Oeiras, Portugal

[3] UCIBIO, Applied Molecular Biosciences Unit, Department of Chemistry, NOVA School of Science and Technology, Universidade NOVA de Lisboa, Caparica, Portugal

[4] Associate Laboratory i4HB, Institute for Health and Bioeconomy, NOVA School of Science and Technology, Universidade NOVA de Lisboa, Caparica, Portugal

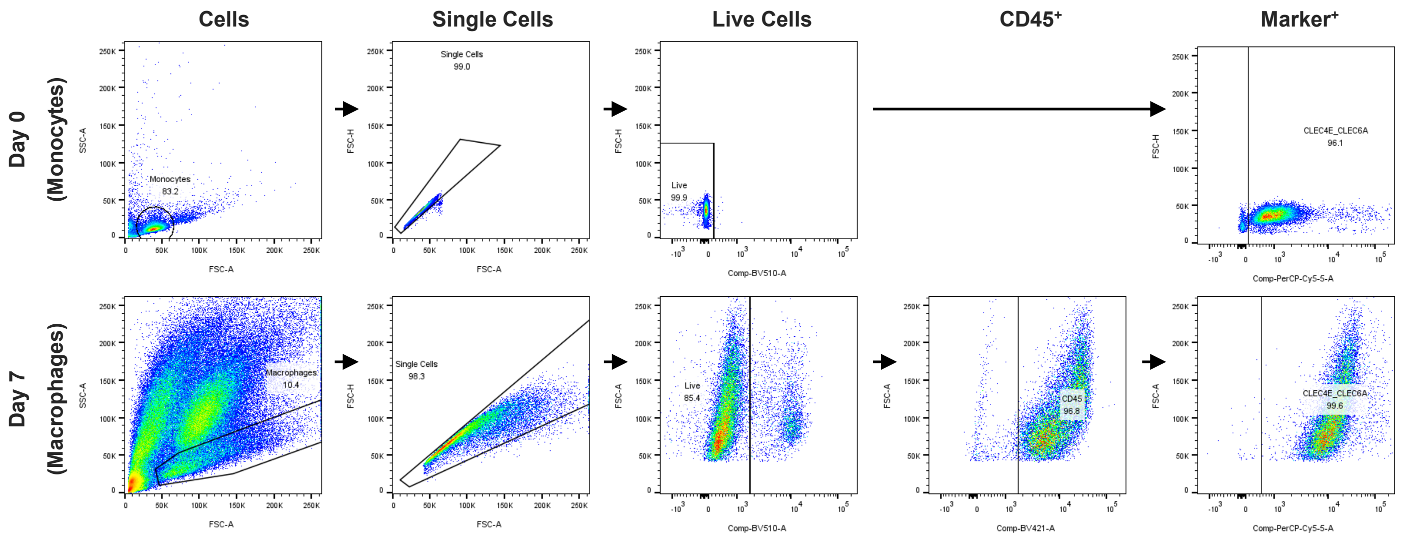


**Supplementary Figure S1.** Gating strategy for flow cytometry analysis of monocytes (Day 0) and macrophages recovered from 3D IS TME cultures (Day 7).

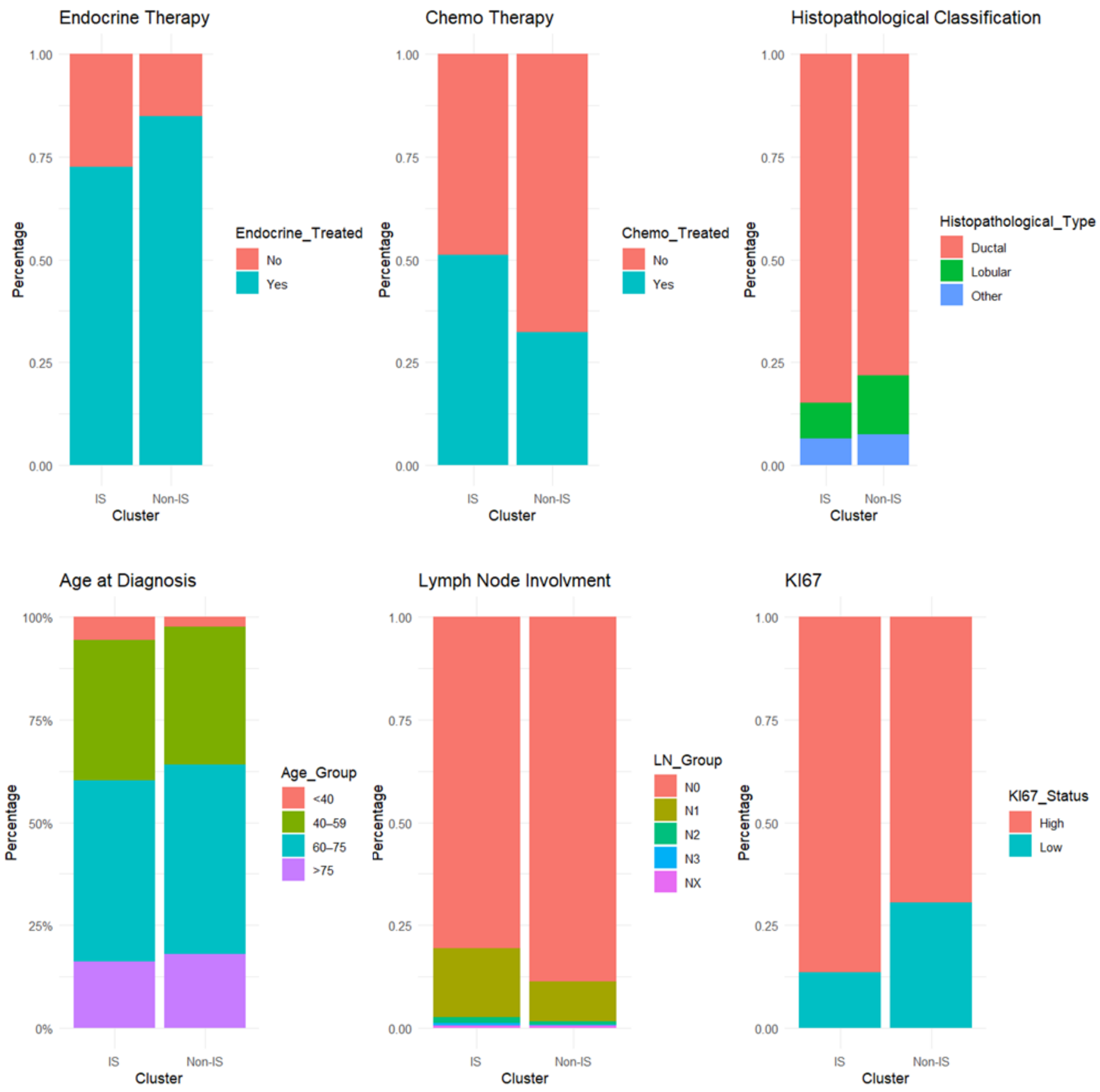


**Supplementary Figure S2.** Analysis of the distribution of clinical parameters in IS and Non-IS clusters.

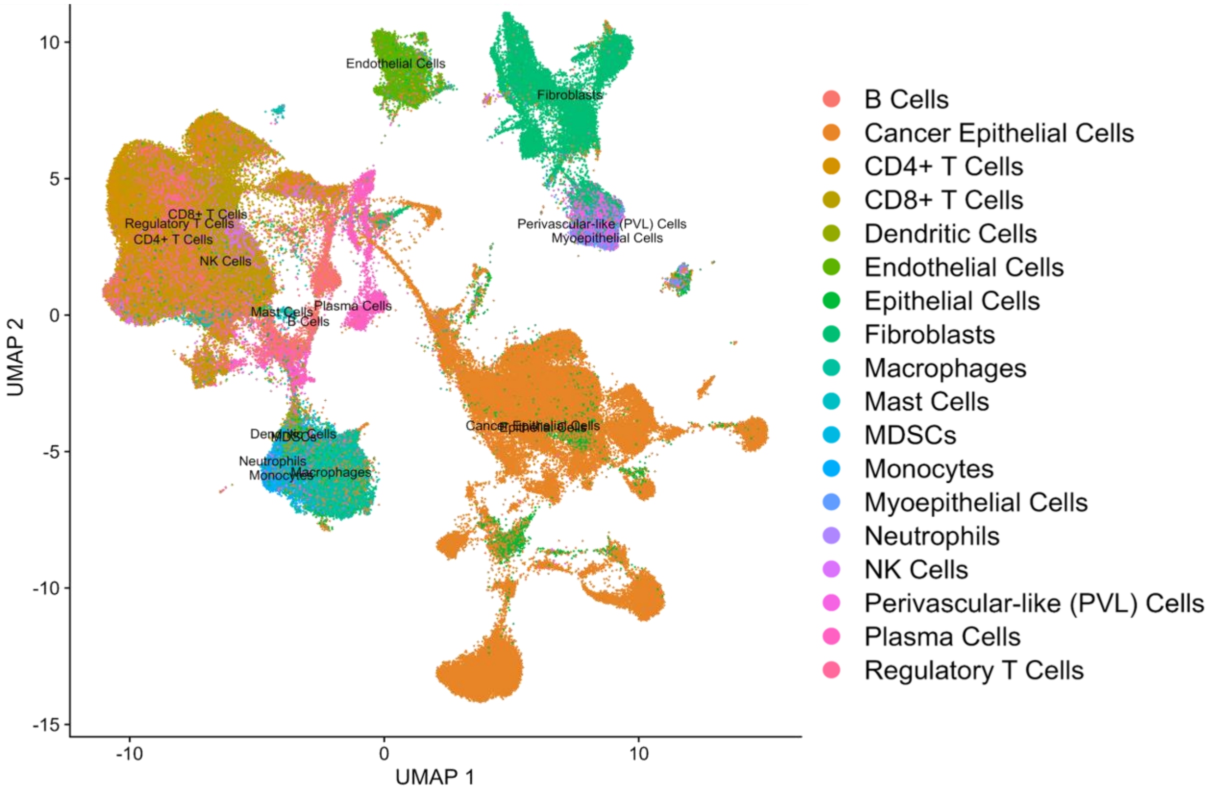


**Supplementary Figure S3.** UMAP of single cell RNA-Seq from the breast tumor atlas of 88 patients (236 363 cells) after QC. Cells are colored by annotated population.

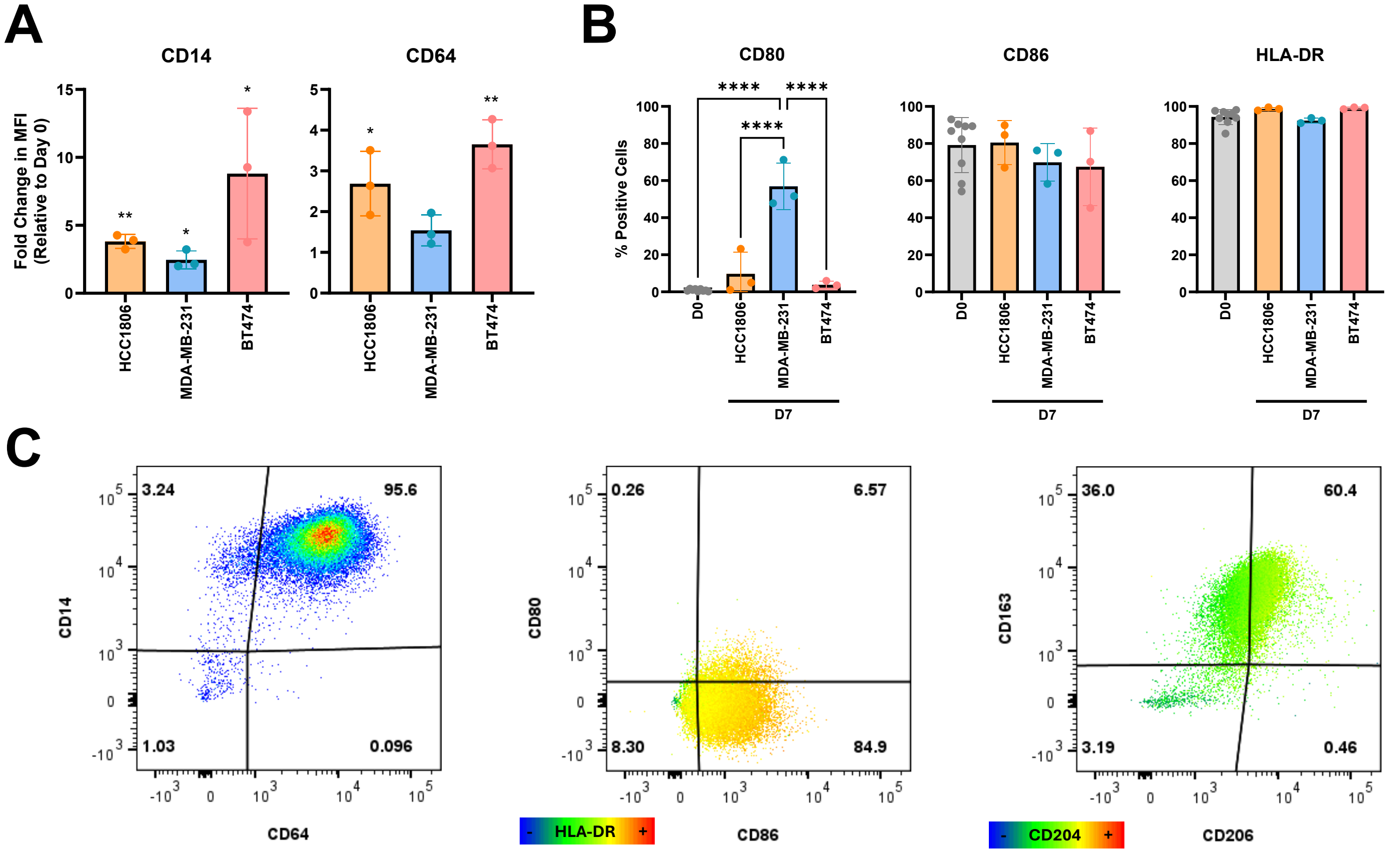


**Supplementary Figure S4.** Fold change in the MFI of CD45+ cells at Day 7 *vs* Day 0 for CD14 and CD64 (A) and percentage of positive cells (relative to live or CD45+ cells) for CD80, CD86 and HLA-DR (B) in monocytes/macrophages in co-culture with HCC1806, MDA-MB-231 and BT474 cells in the 3D IS TME. Representative flow cytometry plots of lineage and polarization markers used for characterization with macrophages from 3D IS-TME cultures with BT474 cell line (C). Statistical significance was determined using one sample ratio t-test (A) or one-way ANOVA with Tukey´s multiple comparisons test (B), with $p < 0.05$ (*), $p < 0.01$ (**), $p < 0.001$ (***) and $p < 0.0001$ (****).

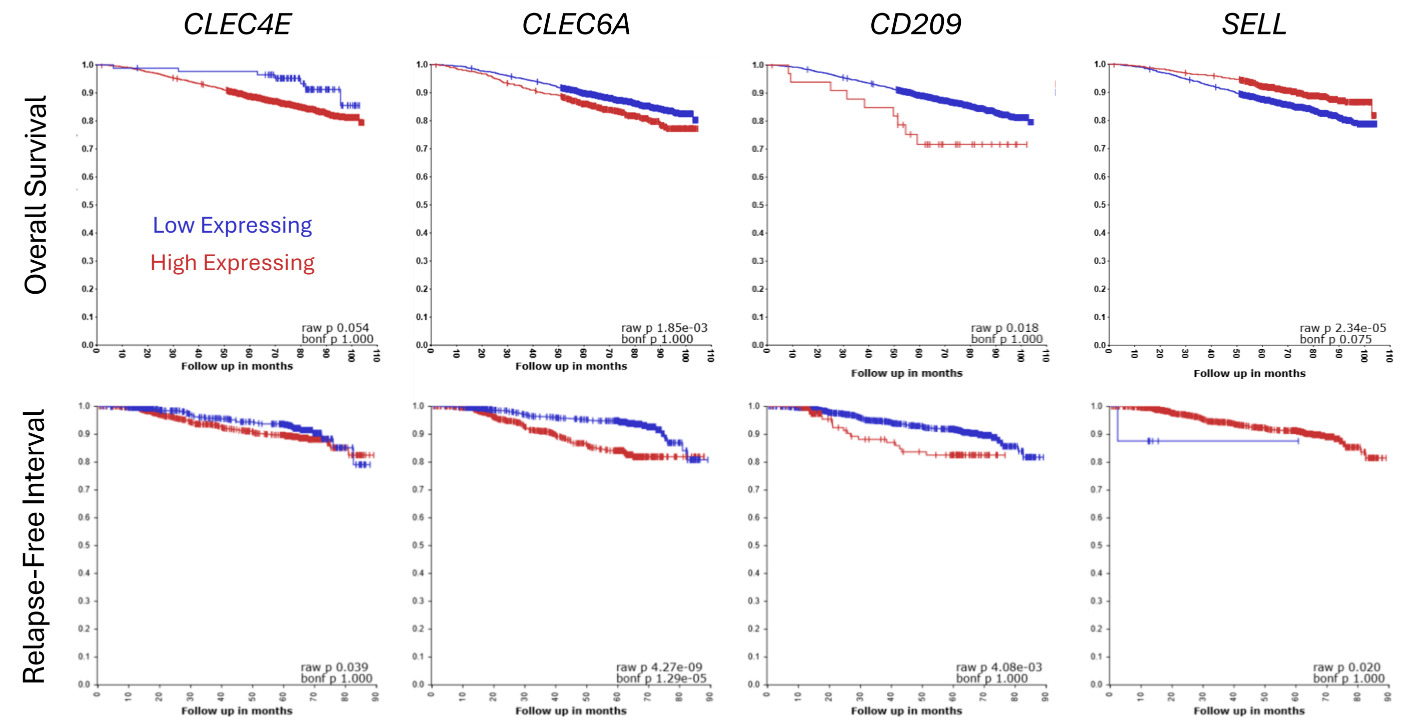


**Supplementary Figure S5.** Kaplan-Meier analysis of overall survival probability and relapse-free interval in selected lectins: *CLEC4E, CLEC6A, CD209* (DC-SIGN) and *SELL*, discriminated by high and low expressing groups.

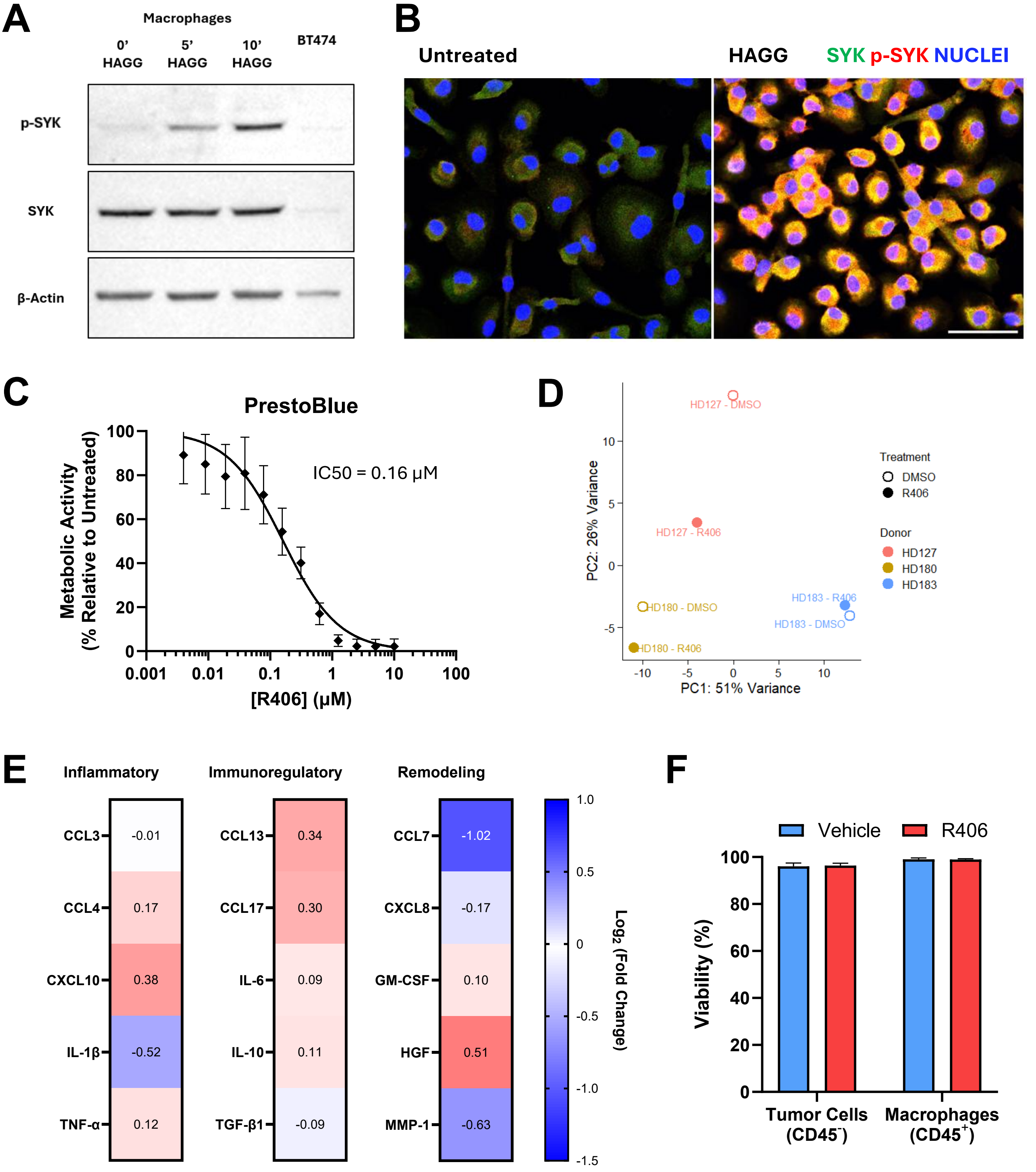


**Supplementary Figure S6**. Validation of SYK activation and R406 tolerability in macrophage cultures. Western blot analysis of p-SYK, total SYK and β-actin in untreated and heat-aggregated IgG (HAGG) stimulated macrophages, confirming inducible SYK phosphorylation (A). Cropped blot; full Length blots are presented in Supplementary Figure S8. Immunofluorescence analysis of untreated and HAGG-stimulated macrophages showing total SYK in green, p-SYK in red and nuclei in blue (DAPI); scale bar = 20 µm (B). Dose–response analysis of R406 tolerability in monocyte-derived macrophages assessed by PrestoBlue metabolic activity assay with IC50 values estimated by nonlinear regression (C). PCA score plot of sorted TAM bulk RNA-seq samples before donor-effect correction, showing that the main source of variation was donor identity rather than treatment condition (D). Heatmap of Olink secretome profiling of vehicle- and R406-treated BT474–monocyte co-cultures with donor-wise median-centered $\log_2$ fold changes (R406/vehicle) (E). Flow cytometry analysis of viability in BT474 tumor cells (CD45-) and macrophages (CD45+) of vehicle- and R406-treated cultures (F).

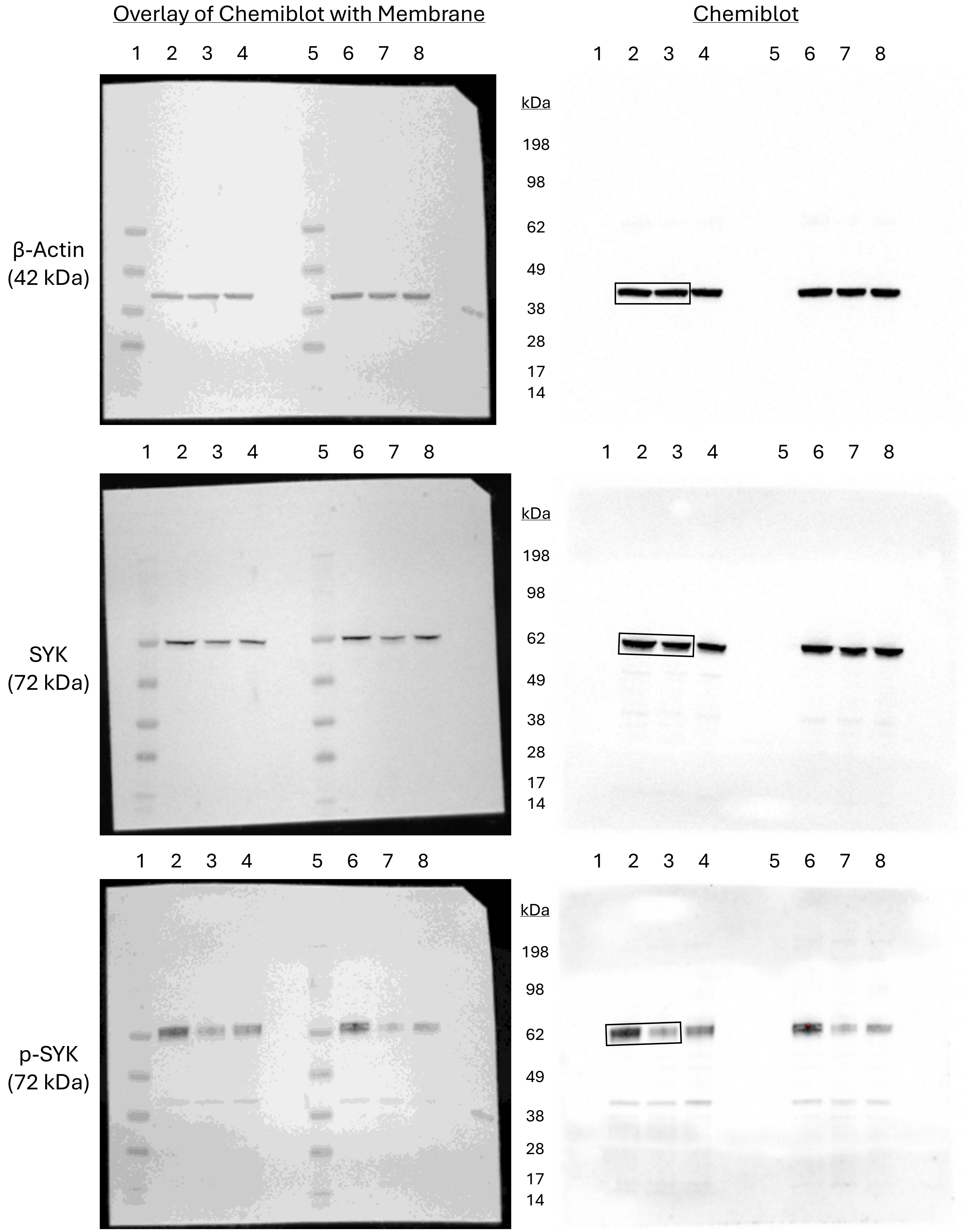


**Supplementary Figure S7**. Full length western blot related to Figure 4A. Lanes: 1) SeeBlue™ Plus2 Pre-stained Protein Standard; 2) Vehicle; 3) R406; 4 to 8) Unrelated Samples. Cropped region indicated by black rectangle.

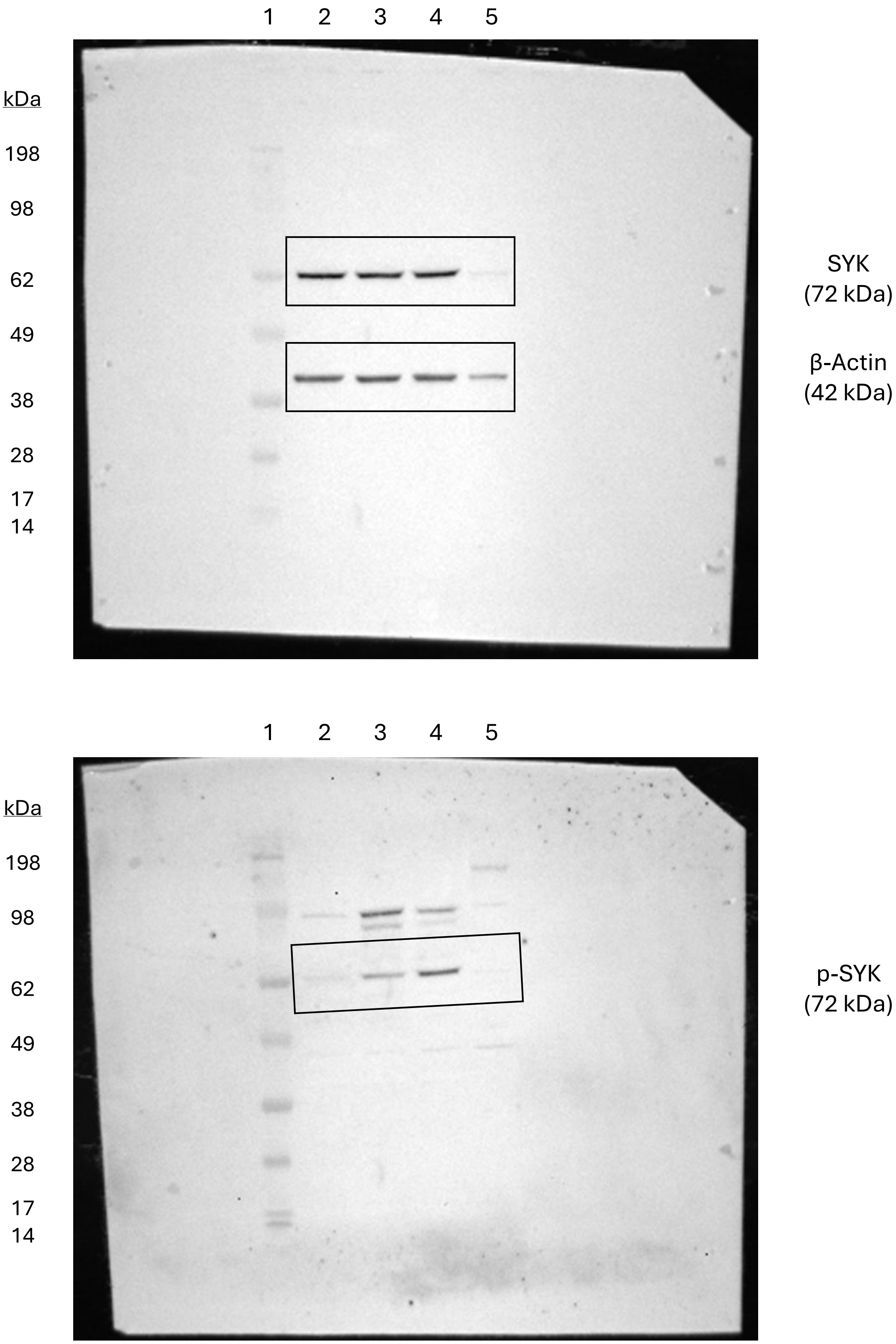


**Supplementary Figure S8**. Full length western blot related to Supplementary Figure S6A. Lanes: 1) SeeBlue™ Plus2 Pre-stained Protein Standard; 2) Macrophages 0’ HAGG; 3) Macrophages 5’ HAGG; 4) Macrophages 10’ HAGG; 5) BT474. Cropped region indicated by black rectangle.

**Supplementary Table S1.** Antibodies used for flow cytometry with final concentration determined by titration.

| Antibody | Brand | Cat # | RRID | [Final] (µg/mL) |
|---|---|---|---|---|
| CD45 – BV421 | BD Biosciences | 563879 | AB_2744402 | 2.5 |
| CD45 – BV605 | BioLegend | 304042 | AB_2562106 | 2.5 |
| CD14 – BV711 | BioLegend | 301838 | AB_2562909 | 7.5 |
| CD64 – PE/Dazzle594 | BioLegend | 305032 | AB_2564186 | 5 |
| CD80 – APC | BioLegend | 305220 | AB_2076147 | 5 |
| CD86 – BV785 | BioLegend | 305441 | AB_2616793 | 5 |
| CD86 – KiraviaBlue520 | BioLegend | 305451 | AB_2892363 | 5 |
| HLA-DR – BV421 | BioLegend | 307636 | AB_2561831 | 2.5 |
| CD163 – APC/Fire750 | BioLegend | 333634 | AB_2734333 | 10 |
| CD206 – AF700 | BioLegend | 321132 | AB_2616869 | 7.5 |
| CD206 - PE | BioLegend | 321106 | AB_571911 | 7.5 |
| CD204 – PE/Cy7 | BioLegend | 371908 | AB_2650772 | 5 |
| CLEC4D – BV786 | BD Biosciences | 744658 | AB_2742398 | 10 |
| CLEC4E – PE/Cy5.5 | Novus Biologicals | NBP1-49311PECY55 | AB_3212754 | 10.6 |
| CLEC6A – PE/Cy5.5 | Novus Biologicals | FAB31141PECY55 | AB_3123816 | 8.3 |
| CD209 – PE/Cy7 | BioLegend | 330114 | AB_10719953 | 2.5 |
| SELL – PE/Cy7 | BD Biosciences | 565535 | AB_2739286 | 10 |
| SIGLEC1 – BV605 | BD Biosciences | 742993 | AB_2741191 | 10 |
| SIGLEC9 – AF488 | R&D Systems | FAB1139G | AB_3645887 | 0.3 |
| CD54 - PE | BD Biosciences | 560971 | AB_2033962 | 10 |
| CD69 – APC | BioLegend | 310910 | AB_314845 | 5 |
| CD72 – BV786 | BD Biosciences | 743799 | AB_2741767 | 10 |
| CD83 – APC | BD Biosciences | 561960 | AB_10894952 | 5 |
| MGL – PE | Miltenyi Biotec | 130-109-582 | AB_2657160 | 5 |

#### Supplementary Methods: HAGG Stimulation and Immunofluorescence

Heat-aggregated IgG (HAGG) was prepared from mouse IgG1 (R&D Systems, #MAB002, RRID:AB_357344) by heating at 63 °C for 30 min and clarified by centrifugation. For SYK activation controls, monocyte-derived macrophage monocultures were stimulated with 200 µg/mL HAGG for 5–10 min and immediately processed for protein extraction or immunofluorescence. For immunofluorescence, macrophages cultured on glass µ-slides (Ibidi) were fixed with 4% PFA/4% sucrose for 20 min at RT, permeabilized and blocked for 30 min in PBS(+/+) containing 0.1% (w/v) Triton X-100 and 0.2% (w/v) fish skin gelatin (FSG). Primary antibodies against phospho-SYK Y525/526 (Cell Signaling Technology, #2710, RRID:AB_2197222) and SYK (Cell Signaling Technology, #80460, RRID:AB_2799953) were diluted in PBS(+/+) with 0.125% FSG and incubated for 2 h at RT followed by overnight incubation at 4 °C. Secondary antibodies anti-mouse Alexa Fluor 594 (Invitrogen, #A11032, RRID:AB_2534091) and anti-rabbit Alexa Fluor 488 (Invitrogen, #A11008, RRID:AB_143165) were used at 1:500. Nuclei were counterstained with 5 µg/mL DAPI for 15 min, and samples were mounted with ProLong Gold Antifade Mountant (Invitrogen). Images were acquired on a Leica Mica microscope.
